# Base editing screens map mutations affecting IFNγ signalling in cancer

**DOI:** 10.1101/2022.03.29.486051

**Authors:** Matthew A. Coelho, Emre Karakoc, Shriram Bhosle, Emanuel Gonçalves, Thomas Burgold, Sarah Cooper, Chiara M. Cattaneo, Vivien Veninga, Sarah Consonni, Cansu Dinçer, Sara F. Vieira, Freddy Gibson, Syd Barthorpe, Claire Hardy, Joel Rein, Mark Thomas, Emile Voest, Andrew Bassett, Mathew J. Garnett

## Abstract

IFNγsignalling underpins host responses to infection, inflammation and anti-tumour immunity. Mutations in the IFNγsignalling pathway cause immunological disorders, haematological malignancies, and resistance to immune checkpoint blockade (ICB) in cancer, however the function of most clinically observed variants remain unknown. Here, we systematically investigate the genetic determinants of IFNγresponse in colorectal cancer cells using CRISPR-Cas9 screens and base editing mutagenesis. Deep mutagenesis of *JAK1* with cytidine and adenine base editors, combined with pathway-wide screens, reveal loss-of-function and gain-of-function mutations with clinical precedence, including causal variants in haematological malignancies and mutations detected in patients refractory to ICB. We functionally validate variants of uncertain significance in primary tumour organoids, where engineering missense mutations in *JAK1* enhanced or reduced sensitivity to autologous tumour-reactive T cells. By classifying > 300 missense variants altering IFNγ pathway activity, we demonstrate the utility of base editing for mutagenesis at scale, and generate a resource to inform genetic diagnosis.

---

Cellular responses to the cytokine interferon *γ* (IFN*γ*) are essential for normal inflammatory responses, but pathway dysfunction and disease can occur through mutation, leading to haematological malignancies and immunological disorders^1, 2^. JAK kinase inhibitors are used to treat myeloproliferative disorders such as polycythaemia vera, and inflammatory disorders such as rheumatoid arthritis and ulcerative colitis^2^, reflecting the central role of JAK-STAT signalling in these diseases. Furthermore, IFN*γ* signalling in cancer cells is a critical aspect of anti-tumour immunity^3, 4^. Clinical resistance to ICB, such as antibody therapies targeting PD-1 and CTLA-4, has been associated with somatic mutation and homozygous inactivation of IFN*γ* pathway components in tumour cells^5–8^, or inactivation of genes involved in antigen processing and presentation (e.g. *B2M*)^9, 10^ that are expressed in response to IFN*γ*. For example, mutations in *JAK1* and JAK2 can confer resistance to ICB^5, 6^. However, such loss-of-function (LOF) mutations in IFN*γ* pathway components are rare, reflecting the limited number of tumour samples sequenced pre- and post-ICB to date^11^, and the apparent absence of convergence (hotspots), which is more common in resistance to small molecule inhibitors^12^. Since somatic mutations in cancer are predominantly single nucleotide changes, which often result in missense mutations with unknown consequence^13, 14^ (i.e. variants of uncertain significance, or VUS), interpreting their functional relevance remains challenging, representing an impediment to diagnosis, patient stratification, and management of drug-resistant disease.

Experimental approaches are instrumental in assessing the functional effects of VUS. This is due to the ability to establish causality between VUS and disease-related phenotypes, as well as a scarcity of clinical datasets (e.g. from sequencing ICB-resistant tumours), and the infrequent occurrence of some variants in patient cohorts. For example, cDNA-based expression of variant alleles can be used^12^, but this is not easily scaled and does not reflect physiological levels of gene expression. Bioinformatic predictions of variant effect are not completely predictive and often discordant^15^. Saturation genome editing (SGE) using CRISPR-mediated introduction of exogenous homology-directed repair (HDR) templates^15^ is challenging to scale to multiple genes, costly, and often limited to cell lines with high levels of HDR and near-haploid genomes, which can restrict its utility for studying VUS in disease-relevant cell models. Another methodology to prospectively assess endogenous gene variant function at scale is base editing^16–20^; a CRISPR-based gene editing technology that employs cytidine^21^ or adenine^22^ deaminases to install C->T or A->G transitions, respectively, achieving high editing efficiencies with minimal generation of DNA insertions and deletions (indels).

In this study, we use CRISPR-Cas9 screening to identify mediators of sensitivity and resistance to IFN*γ* in colorectal adenocarcinoma (CRC), and use cytidine base editors (CBEs) and adenine base editors (ABEs) to perform mutagenesis of the top-scoring genes, thereby systematically mapping LOF and gain-of-function (GOF) variants modulating IFN*γ* pathway activity (Fig. 1a), including VUS associated with diseases such as cancer.

**Figure 1.**
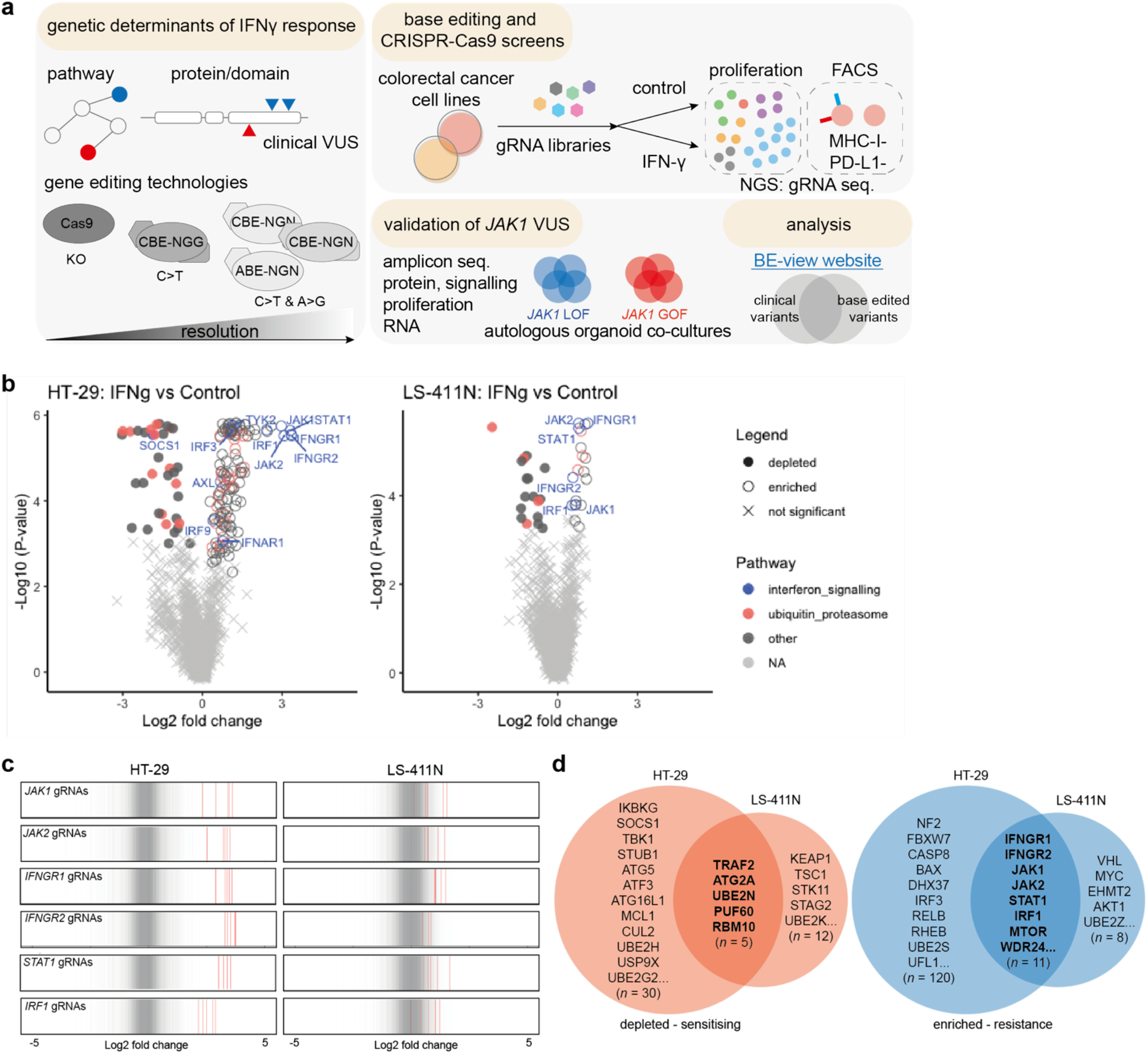
CRISPR-Cas9 screens identify mediators of IFNγ sensitivity and resistance. **a)** Schematic of the integrated CRISPR-Cas9 and base editing screening approaches to identify genetic mediators of sensitivity and resistance to IFN*γ*. Cas9 was used to identify important pathways and genes regulating IFN*γ* response in colorectal cancer cell lines. Multiple base editing mutagenesis screens were used to assess the functional consequence of variants of uncertain significance (VUS) in key regulators. **b)** Gene-level volcano plots of CRISPR-Cas9 screens comparing IFN*γ*-treated to control conditions. Data are the average from two independent screens. **c)** gRNA-level analysis of top resistance genes, representing essential components of the IFN*γ* pathway. **d)** Common and private genes conferring sensitivity and resistance to IFN*γ* in HT-29 and LS-411N CRC cell lines identified from CRISPR-Cas9 screens.

## Results

### CRISPR-Cas9 screens identify mediators of sensitivity and resistance to IFNγ

To systematically evaluate genetic, cell-intrinsic determinants of IFN*γ* signalling, and nominate genes for further investigation, we performed CRISPR-Cas9 screens in two colorectal cancer cell lines, HT-29 and LS-411N (both *BRAF* mutant, and microsatellite stable and microsatellite unstable, respectively) (Fig. 1a). Cas9-expressing derivative cell lines^23^ were transduced with an immuno-oncology focused guide RNA (gRNA) gene knock-out (KO) library, containing 10,595 gRNAs targeting 2,089 genes with a median of five gRNAs per gene (Supplementary Table 1) and selected with cytotoxic doses of IFN*γ*. Screen quality was verified by efficient depletion of gRNAs targeting essential genes^24, 25^ (Supplementary Fig. 1a), and correlation between independent biological screening replicates (Supplementary Fig. 1b).

MAGeCK^26^ (Fig. 1b) and Drug-Z^27^ (Supplementary Fig. 1c) analyses indicated that KO of genes involved in the regulation of IFN*γ* signalling, JAK-STAT signalling, and the downstream transcriptional response, caused the strongest resistance, including *IFNGR1*, *IFNGR2*, *JAK1*, *JAK2*, *STAT1* and *IRF1* (Fig. 1b), each of which had multiple gRNAs with significant enrichment specifically in the presence of IFN*γ* (Fig. 1c and Supplementary Fig. 1d). Changes in gRNA abundance were generally greater for HT-29, reflecting higher sensitivity to IFN*γ* and a faster growth rate than LS-411N (Supplementary Fig. 1e).

Identification of hits common to both cell lines (Fig. 1d) and STRING network analysis^28^ revealed genes centred around IFN*γ* signalling, protein ubiquitination, RNA processing, and mTOR signalling (Supplementary Fig. 1f).

KO of mTOR, AKT1 and WDR24 were significantly associated with resistance to IFN*γ*, whereas negative regulators of mTOR, TSC1 and STK11, were sensitising hits, consistent with the pleiotropic, immunosuppressive effects of rapamycin, and mTOR signalling potentiating IFN*γ* signalling^29^. Gene function enrichment analysis^30^ suggested sensitising and resistance hits were highly enriched for ubiquitin mediated proteolysis and antigen processing pathways^10^ (Fig. 1b, Supplementary Fig. 1g). Inactivation of genes involved in protein degradation such as tumour suppressor genes *KEAP1* and *FBXW7*, have been previously implicated in sensitivity and resistance to cancer immunotherapy, respectively^31, 32^. Interestingly, *FBXW7* was a significant resistant hit in HT-29 but not LS-411N, where *FBXW7* is already mutated^33^. Moreover, sensitising hits included KO of *SOCS1* and *STUB1*^34^, which are negative regulators of IFN*γ* signalling that function through inhibition and proteasomal degradation of JAK1^35^ and IFNGR1 (bioRxiv DOI: 10.1101/2020.07.07.191650). Top-scoring regulators of apoptosis, *CASP8*, *BAX* and *MCL1*, indicated the mode of cell death induced by IFN*γ*, and support the association of *CASP8* mutations with immune evasion in TCGA pan-cancer analyses^9^. Finally, KO of autophagy-related genes enhanced cell death in the presence of IFN*γ* (Fig. 1d; *ATG2A*, *ATG5*, *ATF3* and *ATF16L1*), consistent with autophagy mediating cancer cell resistance to anti-tumour T cells^36^.

Our CRISPR-Cas9 screens identified key nodes of resistance and sensitivity to IFN*γ* in CRC cell lines for further study, with considerable overlap with clinical reports of ICB resistance in patients^5–7^, and genetic screens interrogating cancer immune evasion *in vitro*^10, 31, 36, 37^ and *in vivo*^34, 36, 38^ (Supplementary Discussion, Supplementary Table 1).

### Base editing mutagenesis screening of *JAK1* with BE3-NGG

*JAK1* KO caused robust resistance to IFN*γ* in CRISPR-Cas9 screens, and there is precedence for mutation causing acquired resistance to ICB^5, 6^. Furthermore, *JAK1* somatic mutations in cancer are most frequently missense mutations (58.2 %), with C->T or G->A transition mutations predominating (52.7 %), which can be installed using cytidine base editors (Fig. 2a). Therefore, we set out to use base editing mutagenesis screens to assign functional scores to VUS in JAK1. To deliver large base editor expression constructs and obviate potential toxicity associated with constitutive expression of deaminases, we generated doxycycline-inducible base editor 3 (iBE3)^21^ HT-29 and LS-411N cell lines through a knock-in strategy (Methods). The base editing activity reporter BE-FLARE^39^ estimated base editing efficiency to be ∼40-50% in HT-29 iBE3 (Fig. 2b). Base editing efficiency was considerably lower in LS-411N (Supplementary Fig. 2a), despite both cell lines having similar ploidy (∼3n). Since LS-411N is MSI^33^ with an inactivating mutation in *MLH1*, we tested whether mismatch repair affected base editing^21^ by KO of *MLH1* in HT-29 iBE3 cells (Supplementary Fig. 2b), but found that *MLH1* was dispensable for base editing in this context (Supplementary Fig. 2c).

**Figure 2.**
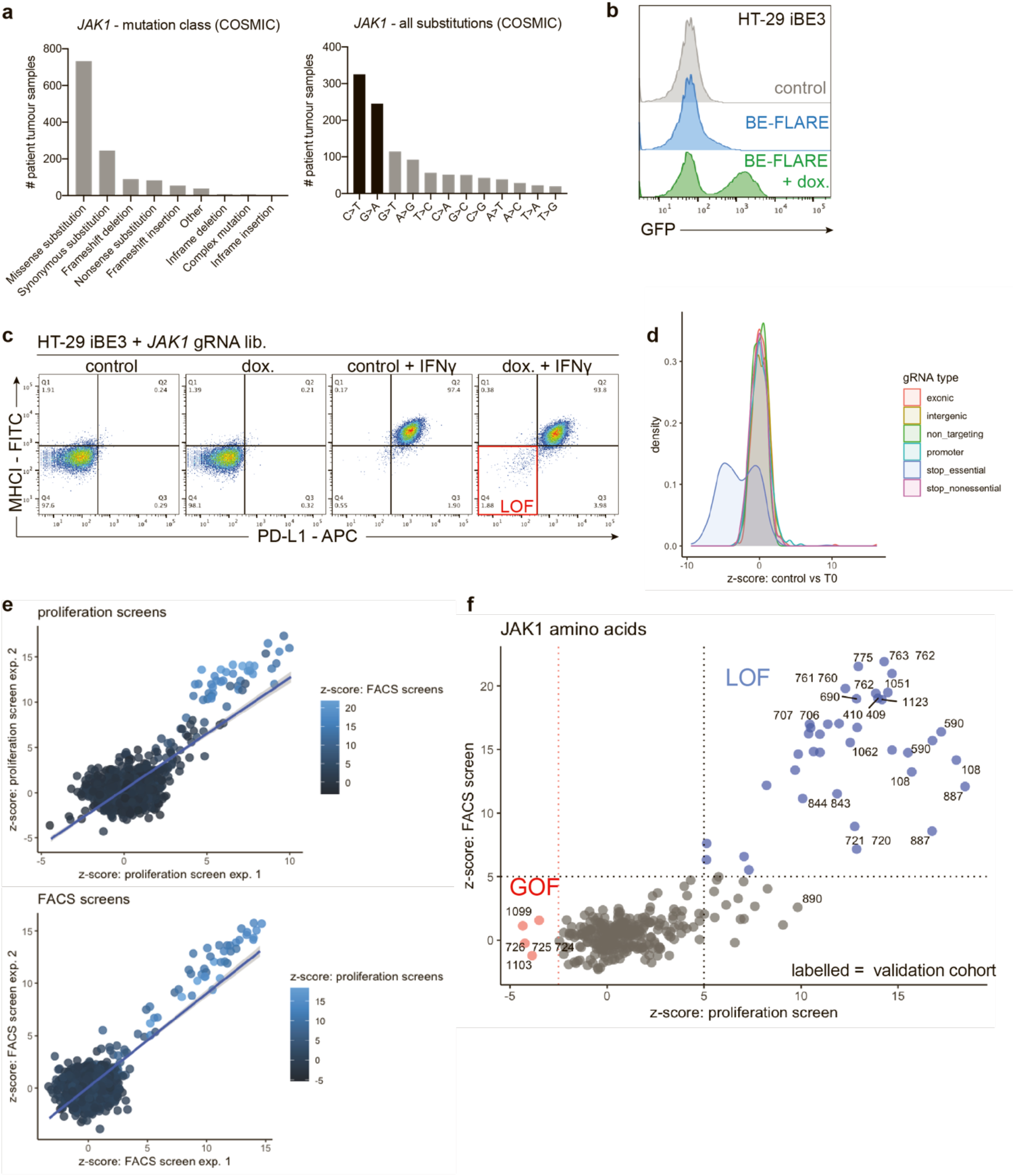
Base editing mutagenesis screening of *JAK1* variants. **a)** COSMIC mutation data from patient tumour samples show *JAK1* mutations in cancer are predominantly C->T and G->A missense variants. **b)** BE-FLARE assessment of base editing efficiency in HT-29 iBE3 cells treated with doxycycline, based on flow cytometry analysis of a BFP (His66) to GFP (Ty66) spectral shift. Data are representative of two independent experiments performed on separate days. **c)** FACS screening assay. After base editing of *JAK1* by the addition of doxycycline, HT-29 iBE3 cells that failed to respond to IFN*γ* after 48 h were selected by FACS, as determined by lack of induction of MHC-I and PD-L1 expression. Data are representative of two independent experiments performed on separate days. **d)** Proliferation screening assay. gRNA depletion or enrichment is indicated by z-score, comparing control arm to T0 (time 0) control sample. Base editing gRNAs designed to introduce stop codons in essential genes in HT-29 iBE3 cells are depleted. **e)** Correlation between screening replicates and different assays. z-scores for gRNAs targeting *JAK1* were compared between replicates and alternative screening assays, with each replicate representing an independent screen performed on a separate day. The shaded line area represents the 95 % confidence interval. **f)** Identification of LOF and GOF alleles in *JAK1* protein affecting sensitivity to IFN*γ*. z-scores for the base editing screens using FACS vs proliferation were plotted to robustly select potential LOF (blue) and GOF (red) *JAK1* variants. Labelling illustrates amino acid positions that were selected for further validation.

Using a pooled library of 2,000 gRNAs, we tiled *JAK1* in HT-29 iBE3 cells with 665 exon-targeting gRNAs and gRNAs targeting *JAK1* promoter regions, non-targeting (NT), intergenic targeting, and controls gRNAs designed to introduce stop codons in 72 essential and 28 non-essential genes (Supplementary Table 2). We adopted two screening approaches; a long-term proliferation screen, and a short-term flow cytometry-based assay, based on MHC-I and PD-L1 induction with IFN*γ* (Fig. 2c). gRNAs predicted to cause stop codons within essential genes were significantly depleted (Fig. 2d), achieving recovery of known essential genes in both screens (AUC = 0.65; Supplementary Fig. 2d). There was no relationship between gRNA functional scores and the number of off-target sites^40^ (Supplementary Fig. 2e), however, the gRNA Rule Set 2 score^41^ (*P* = 9.0x10^-4^; Supplementary Fig. 2f), or considering the immediate sequence context of the target cytidine^21^ (Supplementary Fig. 2g), was somewhat predictive of gRNA performance^19, 20^. Correlation between independent replicates (Fig. 2e; proliferation R^2^adj. 0.58; FACS R^2^adj. 0.68) and the proliferation and FACS screens (R^2^adj. 0.68), was driven by highly enriched gRNAs after positive selection with IFN*γ*, representing candidate *JAK1* LOF variants. As GOF variants were rare, we could only practically sort for *JAK1* LOF cells by FACS, and so only recovered LOF gRNAs in the FACS screen (Supplementary Fig. 3a). We selected 24 gRNAs for downstream validation studies, representing 15 LOF and 5 GOF unique variants, mostly predicted to generate missense variants with clinical precedence in cancer (Fig. 2f; validation cohort). In addition, we included *JAK1* Glu890 gRNA, which was unusual as it scored in the proliferation screens but not the FACS screens (Fig. 2f), and the Trp690* gRNA as a control; predicted to generate a nonsense mutation observed in a CRC patient that failed to respond to ICB^6^.

### Base editing mutagenesis of the IFNγ pathway

Having established a robust base editing system, to achieve a more comprehensive overview of functional missense mutations in the IFN*γ* pathway, we expanded our base editor mutagenesis screens to include top hits of our CRISPR-Cas9 screen using HT-29 iBE3-NGG (Fig. 1b). We tiled *JAK1*, *JAK2*, *IFNGR1*, *IFNGR2*, *STAT1*, *IRF1*, *B2M* and *SOCS1* with 4,608 gRNAs, including the previous *JAK1* gRNAs to serve as internal controls (Fig. 3a). *B2M* was included because of its role in MHC-I presentation and anti-tumour immunity, but it was not a hit in our initial IFN*γ* survival screens as *B2M* variants should not have an effect on cell proliferation *in vitro*.

**Figure 3.**
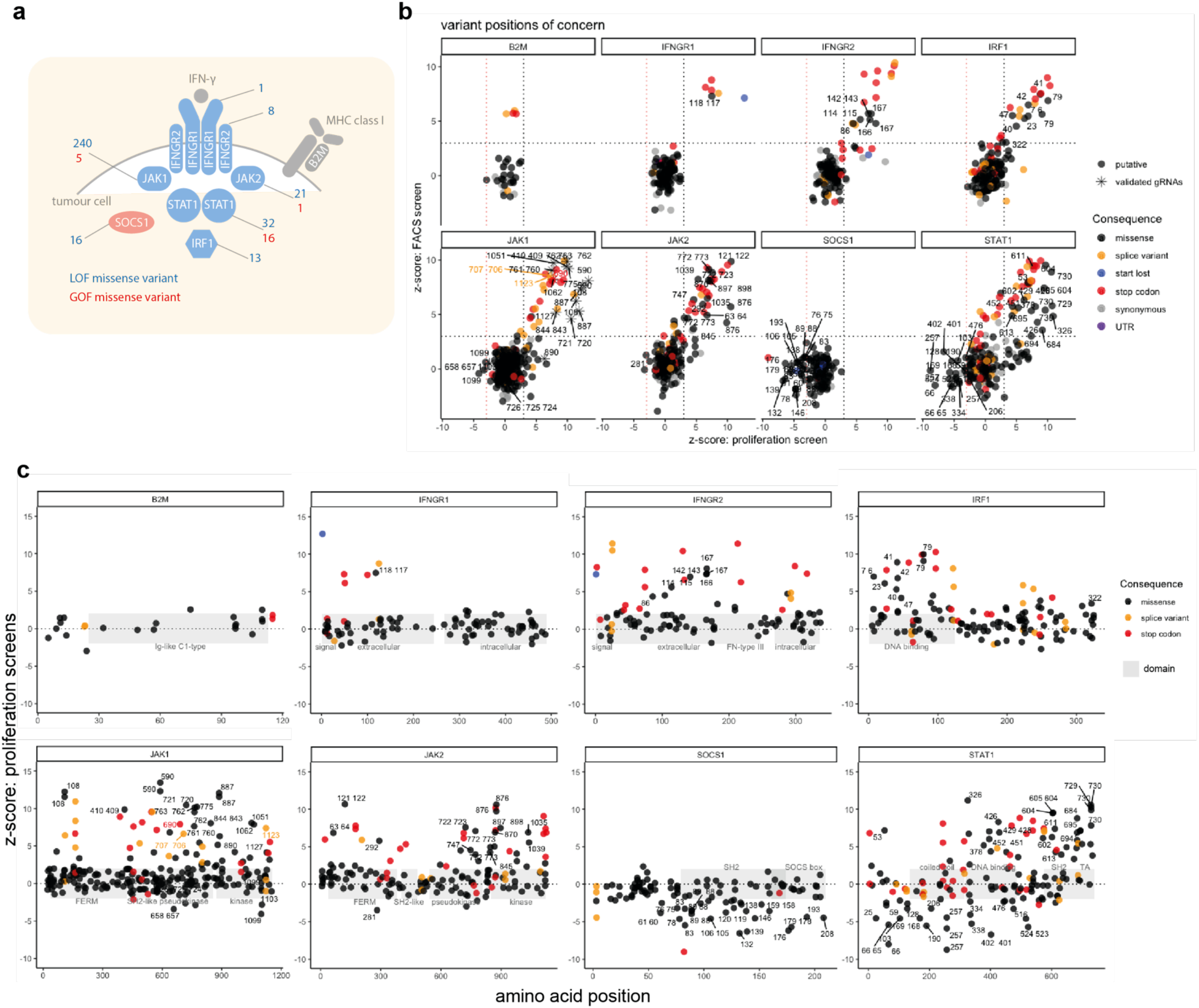
Base editing mutagenesis of the IFNγ pathway. **a)** Schematic of the key mediators of IFN*γ* signalling investigated in base editing screens. Depicted are top hits from our CRISPR-Cas9 screens to determine modulators of sensitivity to IFN*γ*; positive mediators are in blue and negative regulators are in red. LOF and GOF missense variants revealed from all base editing screens are indicated. **b)** Base editor mutagenesis of core IFN*γ* pathway components reveals GOF and LOF missense mutations. The average FACS screen score is plotted against the average proliferation screen score for each gene. Positions of validated *JAK1* gRNAs and amino acid positions with missense LOF or GOF effect are labelled. **c)** Base editing reveals the position of functional domains. Schematics of the domain architecture of proteins in the IFN*γ* pathway tiled with base editing gRNAs, with the distribution of GOF and LOF amino acid positions labelled.

Proliferation and FACS screens were significantly correlated (R^2^adj. 0.42), as were independent replicate screens (proliferation R^2^adj. 0.37; FACS R^2^adj. 0.34; Supplementary Fig. 3b), each displaying a high level of enrichment of gRNAs predicted to introduce splice variants, stop codons and start-lost mutations (Fig. 3b). Once again, *JAK1* Glu890 gRNA was enriched in the proliferation screen, but not in the FACS screen. Such behaviour was rare for most proteins except for the transcription factor *STAT1*, where a cluster of LOF missense mutations was enriched only in the proliferation screen (Fig. 3b), possibly indicating separation-of-function mutants. Encouragingly, we recovered validated gRNAs targeting *JAK1* in this larger screen (Supplementary Table 3, and later sections). In addition to protein truncating mutations, we used *JAK1* LOF and GOF gRNAs from our validation cohort as a benchmark for setting the thresholds to call high-confidence functional missense variants in the IFN*γ* pathway (Fig. 3b).

Due to its short gene length, we only recovered highly enriched gRNAs predicted to install splice site or stop codon variants in *B2M*, and these only scored in the FACS screen, as expected. For the negative regulator of IFN*γ* signalling, *SOCS1*, LOF mutations were significantly depleted. Editing of *JAK1*, *JAK2*, *IFNGR1*, *IFNGR2* and *IRF1* predominantly gave rise to LOF missense mutations, but *STAT1* was a notable outlier as it displayed a high proportion of GOF mutations (Fig. 3b). 66.7 % of STAT1 LOF missense variants were clustered around the SH2 and transactivation domains, compared to 6.7 % of nonsense and splice LOF mutations (Fig. 3c). Conversely, 55.6 % of STAT1 GOF mutations were within coiled-coil and DNA-binding domains, consistent with previous reports of GOF mutations in patients with chronic mucocutaneous candidiasis (CMC)^42^. LOF missense mutations in IRF1 were enriched in the DNA binding domain (88.9 %), whereas SOCS1 LOF missense variants were enriched in the SOCS box and SH2 domains (84.2 %) or within the JAK kinase inhibitory region^35^ (SOCS1 His61Tyr), demonstrating that base editing can highlight functional protein domains.

### Comparison of base editing technologies for mutagenesis screening

Analysis of amino acid mutations predicted from gRNA sequences suggested the BE3-NGG library targeted approximately 21.4 % amino acids in JAK1. To improve the saturation of mutagenesis achievable with base editing, we employed a Cas9 variant with a relaxed NGN PAM requirement^43^, generating BE3.9max-NGN^20, 44^. Secondly, we sought to increase product purity by using a YE1-BE4max-NGN architecture that reduces non-C->T outcomes^44, 45^, reduces Cas9-independent off-target editing, and improves editing precision by employing an engineered deaminase (YE1) with a narrower editing window^46, 47^. Finally, we employed an adenine base editor^22^ (ABE8e-NGN)^48^, to incorporate a wider variety of amino acid substitutions than can be achieved by C->T transitions alone.

Using our panel of iBE3-NGG, iBE4max-YE1-NGN, iBE3.9max-NGN and iABE8e-NGN HT-29 base editor cell lines, we re-screened *JAK1* with a library of 3,953 gRNAs (Fig. 4a, 4b), consisting of the 2,000 gRNA *JAK1* gRNA iBE3-NGG library and an additional 1,953 NGT, NGC and NGA gRNAs targeting *JAK1* exons (Supplementary Table 4). For comparison, we included *JAK1* screening data from pathway-wide base editing screens. For NGN base editors, we detected significantly enriched gRNAs utilising all four PAMs (Supplementary Fig. 4a). Missense variants displayed the most heterogeneous phenotype (Supplementary Fig. 4b). ABE cannot introduce stop codons, but predicted splice variants in JAK1, which could be introduced with both CBE and ABE, were significantly enriched over NT control gRNAs in all screens (Supplementary Fig. 4b). Given the PAM utility and editing windows of each base editor, we predicted non-synonymous amino acid mutation coverage of *JAK1* was improved to approximately 39.6 % for BE4max-YE1-NGN, 50.8 % for BE3.9max-NGN, 64.9 % for ABE8e-NGN, and 85.1 % when combining cytidine and adenine NGN mutagenesis. However, we cannot guarantee the editing efficiency of all gRNAs, so the absence of a significant score cannot be used as evidence for the lack of function of an amino acid position.

**Figure 4.**
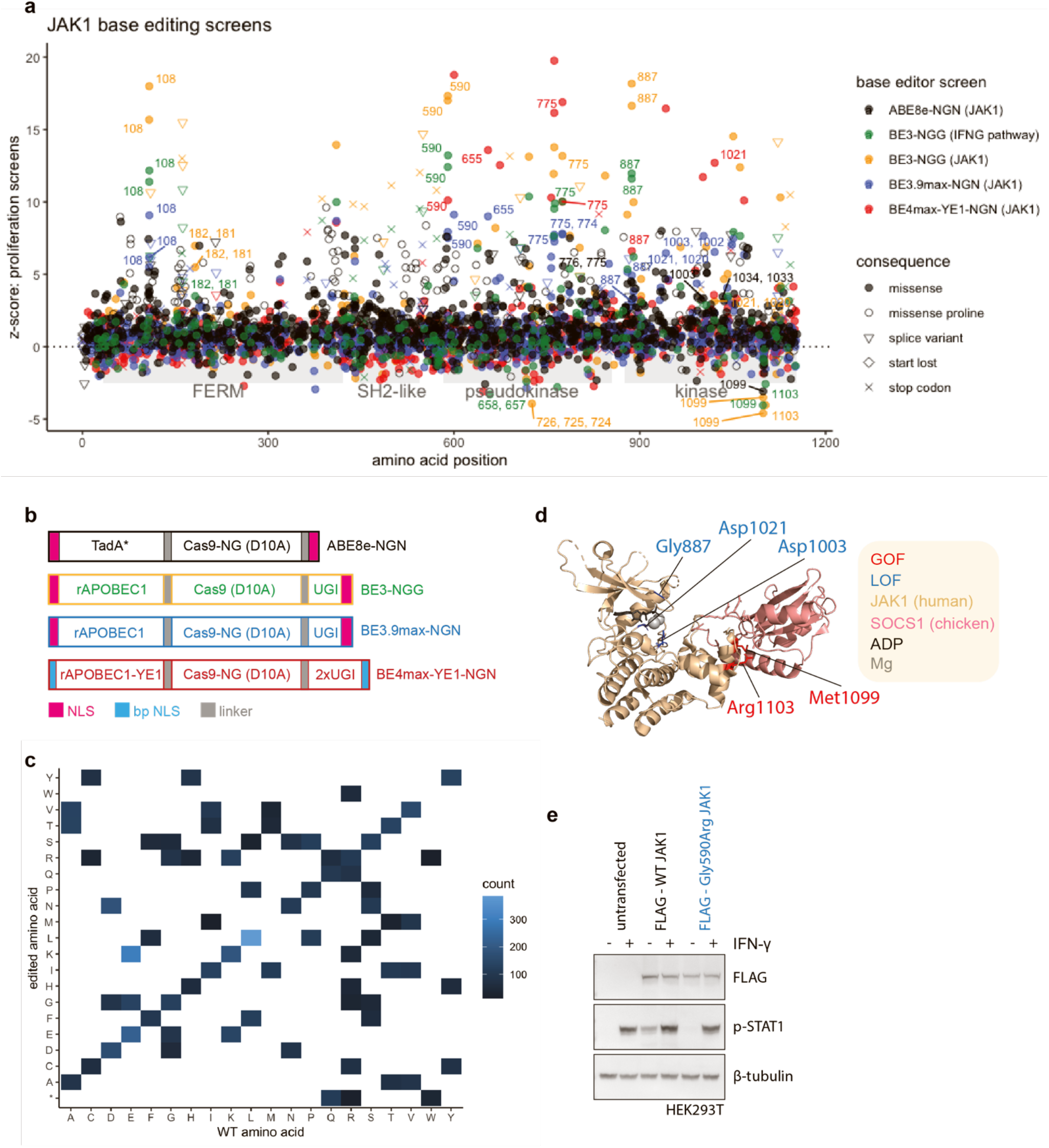
Base editing reveals *JAK1* LOF and GOF variants with clinical precedence. **a)** Functional variant map of JAK1. z-scores from base editing proliferation screens are plotted for each gRNA across *JAK1* protein domains. gRNAs producing candidate LOF and GOF positions referred to in the text are labelled with the predicted edited amino acid positions. Synonymous variants are not shown. Screen z-scores are calculated independently for each base editor and plotted together for comparison. **b)** Schematic of base editor architectures used in screening experiments. Bp NLS; bipartite nuclear localisation sequence. **c)** Heatmap showing the frequency of predicted amino acid substitutions in *JAK1* when merging CBE and ABE-NGN base editing screens. **d)** Structural insight into the mechanism of action of *JAK1* LOF and GOF mutations. Crystal structure (6C7Y) shows catalytic LOF mutations (blue) proximal to the ATP/ADP binding pocket in the kinase domain, and GOF mutations (red) in the binding interface with the negative regulator SOCS1. **e)** Western blotting analysis of HEK293T cells following overexpression of FLAG-tagged WT or Gly590Arg mutant JAK1, with or without IFN*γ* stimulation for 1 h. Reduced p-STAT1 signalling is independently replicated in Supplementary Fig. 5d.

When combined, CBE and ABE editors can achieve substitutions of all 20 amino acids to at least two alternative amino acids (Fig. 4c). Notably, substitution of amino acids with disparate chemical properties achieved larger average effect sizes, such as Gly->Glu (CBE) and Phe->Ser (ABE) (Supplementary Fig. 4c). One exceptional outlier specific to ABE editing was Leu->Pro missense mutations, which were significantly enriched in LOF mutations over other missense variants (*P* = 2.2x10^-16^), presumably due to the uniquely restricted *ϕ*and *ψ*peptide bond angles available to proline (Fig. 4a, Supplementary Fig. 4c). Comparison of functional scores to *in silico* predictions of variant effect (SIFT, PolyPhen and BLOSUM62) demonstrated imperfect predictions in each case (Supplementary Fig. 4d), implying high-throughput experimentation is often required to complement bioinformatic prediction of variant effect^49^.

Functional comparison of BE3 and BE4max-YE1 editing of *JAK1* confirmed the narrower editing profile of the YE1 engineered deaminase (Supplementary Fig. 5a), but we observed a lower editing efficiency for BE4max-YE1-NGN compared to the WT deaminase (Supplementary Fig. 5b), consistent with a reduced number of significant missense, splice and stop codon variants compared with alternative NGN base editor architectures (Supplementary Fig. 4a, 4b). As expected, functional gRNAs present in both BE3 and BE4max-YE1 screens had target cytosines within the YE1 5-7 activity window (e.g. Asp775Asn gRNA 908510028), whereas out-of-window targeting gRNAs were not enriched in the BE4max-YE1 screens (e.g. Trp690* gRNA 908510274; Supplementary Fig. 5a, 5c).

**Figure 5.**
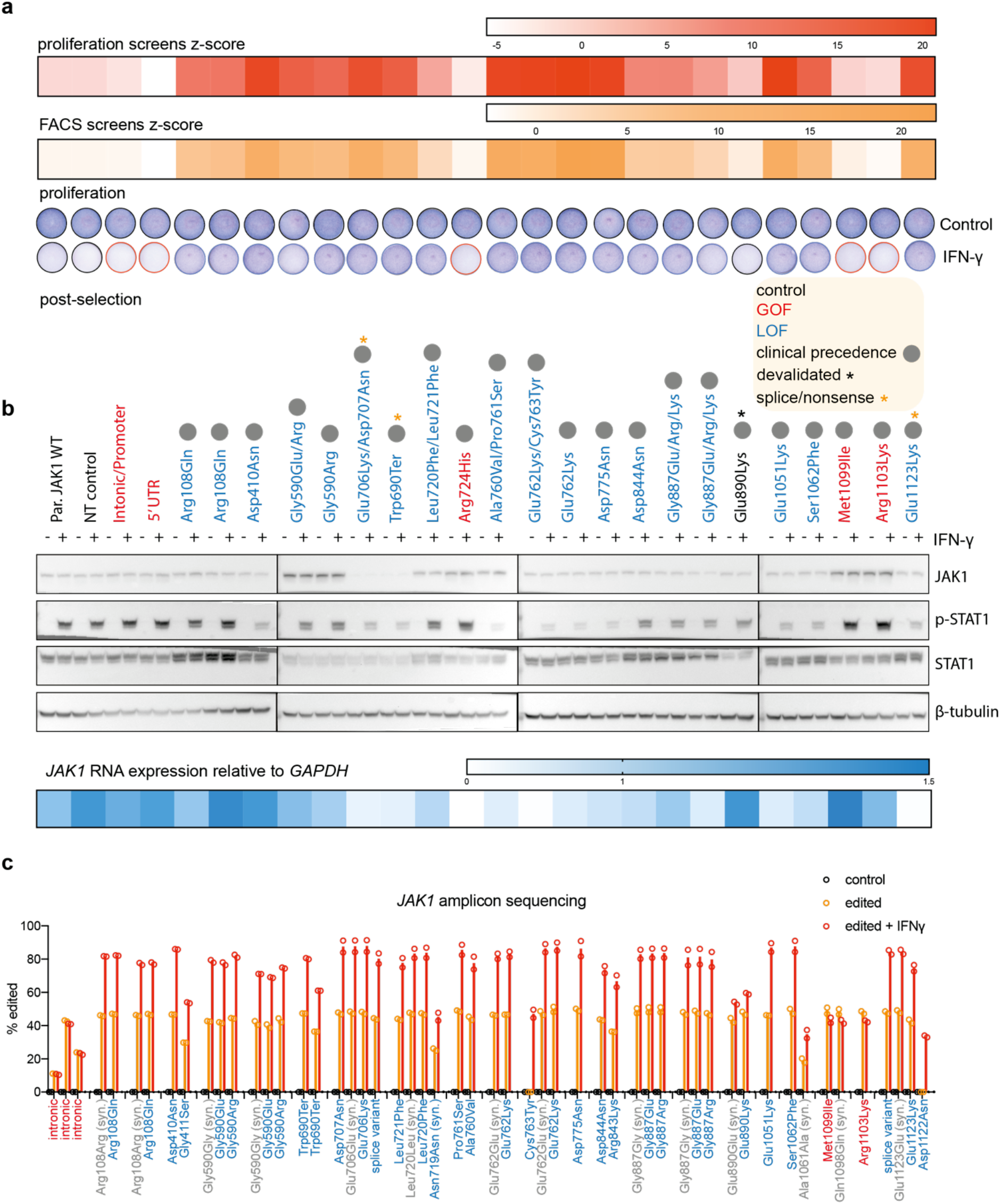
Functional validation of variants conferring altered sensitivity to IFNγ. **a)** Functional validation of base editing gRNAs targeting *JAK1* in HT-29 iBE3 cells. Proliferation assay: Giemsa stain following growth in the presence or absence of IFN*γ*. All data are representative of two independent experiments performed on separate days. Base editing screen z-scores for each gRNA are provided for comparison. See also Supplementary Fig. 4e. **b)** Western blotting analysis of *JAK1* expression and p-STAT1 signalling of corresponding *JAK1* variants was performed on cells stimulated with IFN*γ* for 1 h, after selection in IFN*γ* for LOF variants. RNA expression: qPCR analysis of *JAK1* RNA expression relative to *GAPDH* 72 h after base editing. All data are representative of two independent experiments performed on separate days. See also Supplementary Fig. 4e. **c)** Deep sequencing of *JAK1* reveals the DNA editing profile of base editor gRNAs. Editing variant allele frequency for LOF and GOF gRNAs within the validation cohort measured by NGS of amplicons in control cells, base edited cells, or base edited cells with selection with IFN*γ* for 6 d. Different editing outcomes are grouped by gRNA. Syn; synonymous. Data represent the mean of two independent experiments performed on separate days.

### Deep mutagenesis of *JAK1* reveals LOF and GOF variants with clinical precedence

To aid interpretation of our mutagenesis screens, we compiled a database of variants from COSMIC^13^, TCGA, ClinVar^50^, gnomADv3.1^51^, literature on functional *JAK1* mutations^1^, post-translational modifications^52^, and clinical data from patients receiving ICB where cancer exome sequencing data is publicly available^6, 8, 52–57^, and aligned this with predicted *JAK1* variants. This analysis revealed GOF variants in the *JAK1* pseudokinase domain with clinical precedence in cancer. gRNAs targeting position Arg724 were significantly depleted with IFN*γ* (Fig. 4a). The base edited variant, Arg724His, has been implicated in activating *JAK1* signalling in acute lymphoblastic leukaemia through dysregulating intramolecular inhibition of the kinase domain^1^. Another GOF position, *JAK1* Val658, is mutated in acute myeloid leukaemia (AML); this residue is structurally analogous to JAK2 Val617, which is commonly mutated in polycythaemia vera^1, 2^. CBE and ABE screens converged on a cluster of GOF variants in the C-terminus of the kinase domain (Met1099, Arg1103) in a known protein-protein interaction motif for SOCS1^35^ (Fig. 4a); a significant negative regulator in our CRISPR-Cas9 screens. These variants presumably disrupt this interaction, increasing *JAK1* protein abundance and activity (Fig. 4d). Indeed, amplification of SOCS1 has been found in patients that failed to respond to ICB^7^, implying this regulatory mechanism is of clinical relevance.

LOF positions included Gly887 (Fig. 4a), which is within the kinase active site, with the crystal structure^35^ suggesting mutation of this residue would negatively affect Mg^2+^ and ATP/ADP coordination (Fig. 4d). Other LOF mutations involving kinase catalytic residues included Asp1003 (proton acceptor), and Asp1021 (within the DFG motif), which were detected with increased (NGN) saturation (Fig. 4a). ABE screens were more likely to detect sites of post-translational modification due to the ability to modify tyrosine, threonine and serine (phosphorylated) and lysine (ubiquitinated), revealing Tyr993, and the known activating Tyr1034 phosphosite as candidate LOF positions (Fig. 4a), and Lys267 as a GOF site. LOF and GOF variants made with CBE were more likely to be clinically apparent than ABE variants; with 91 % of CBE functional variants identified occurring at residues with precedence of mutation in humans and in cancer genomes^1, 6, 8, 13, 50–57^, 42 % of which were predicted to be recapitulated with CBE (vs 7 % for ABE), perhaps reflecting the APOBEC deamination signature in cancer^14^.

Candidate LOF mutations Gly655Asp^58^, Gly182Glu^54^ and Gly590Glu^54^ (Fig. 4a) were all VUS detected in patients that failed to respond to ICB. In addition, *JAK1* Asp775Asn has been independently verified as a LOF variant in melanoma^6^. To verify the potential functional significance of the Gly590 clinical variant, we transiently expressed FLAG-tagged WT *JAK1* or Gly590Arg *JAK1* in HEK293T cells. As endogenous *JAK1* is also present, the cells responded normally to IFN*γ* as measured by phosphorylation of STAT1. WT *JAK1* overexpression resulted in pSTAT1 signal even in the absence of IFN*γ*, and supraphysiologic stimulation with IFN*γ*, whereas the *JAK1* Gly590Arg mutant failed to induce STAT1 phosphorylation to the same extent in either context (Fig. 4e), verifying this clinical VUS as a *bona fide* LOF mutation.

In sum, these data demonstrate that mutagenesis screens utilising multiple different base editor architectures and deaminases can be effectively integrated to assign function to clinically relevant VUS over an entire protein. We assign function to > 200 missense variants in *JAK1* affecting protein function, with multiple predicted mechanisms of action, including conformational protein-protein interaction interfaces (SOCS1-JAK1), catalytic residues (ATP coordination), post-translational modifications, structural variants (missense proline), and complex intra-molecular interaction interfaces (JAK1 pseudokinase-kinase domain). Across all base editing screens, we have identified 358 LOF and 22 GOF missense mutations altering IFN*γ* signalling (Fig. 3a), many of which had previously unknown function (Supplementary Table 5).

### Functional validation of variants conferring altered sensitivity to IFNγ

We set out to functionally validate 24 gRNAs comprising our *JAK1* validation cohort (Fig. 2f) in an arrayed format, with multiple assays assessing cell proliferation, signalling, protein expression and RNA expression (Fig. 5a and Fig. 5b). This analysis was germane to screening results from multiple base editing modalities, due to their convergence on *JAK1* residues within the validation cohort (e.g. Arg108, Gly590, Asp775, Gly887, Met1099; Figure 4a and Supplementary Table 5). The growth of HT-29 iBE3 cells with engineered *JAK1* variants in the presence of IFN*γ* tracked with screen z-scores, with GOF variants having no survival benefit and LOF variants having robust resistance to IFN*γ*, relative to controls (Fig. 5a, Supplementary Fig. 5d). Base editing screening gRNAs were validated, with the possible exception of *JAK1* Glu890, which scored poorly in the FACS screen (Fig. 2f), highlighting the value of implementing two screening assays.

Many of the candidate LOF variants had reduced levels of pSTAT1 induction, whereas GOF variants had enhanced levels of pSTAT1 (Fig. 5b, Supplementary Fig. 5d). Met1099 and Arg1103 GOF variants had increased levels of *JAK1* protein and JAK-STAT signalling, consistent with disruption of the SOCS1 binding interface and reduced E3 ubiquitin ligase-mediated destruction^35^. Surprisingly, the Gly590 LOF variants also had elevated levels of *JAK1* protein (Fig. 5b, Supplementary Fig. 5d), despite reduced sensitivity to IFN*γ* in terms of cell proliferation and signalling. We speculated that increased *JAK1* Gly590Arg protein could also be attributable to altered binding to SOCS1, however, we did not observe any change in binding in co-immunoprecipitation experiments (Supplementary Fig. 5e). *JAK1* 706/707 gRNA targets a splice region and had severely reduced *JAK1* protein expression similar to the clinical Trp690* nonsense control (Fig. 5b). The Glu1123 splice variant reduced *JAK1* RNA abundance to levels comparable to the Trp690* nonsense control, which we presumed was targeted for nonsense mediated decay. However, basal *JAK1* variant RNA expression levels were generally only modestly affected and RNA expression was not entirely indicative of *JAK1* protein levels, consistent with the high level of post-translational control of JAK1^35^.

Finally, we performed amplicon sequencing of the endogenous *JAK1* loci to unambiguously assign base edited genotypes (Fig. 5c). This analysis confirmed predictions of base editing outcomes, detecting C->T editing focused within the BE3 activity window (∼4-9 relative to the PAM at position 21-23), with the minority of gRNAs (22.7 %) exhibiting lower frequency edits upstream or downstream, ranging between positions -11 to 11 (Supplementary Fig. 6). Collectively, this resulted in two unanticipated coding mutations from the validation cohort (JAK1 Asp1122Asn and Gly590Glu) caused by editing at protospacer positions 2, 3 and 11. LOF variants were enriched in the presence of IFN*γ* without exception (JAK1 Glu890 was modestly enriched), demonstrating functionality. Co-enrichment of LOF variants with synonymous mutations (63.6 % of gRNAs) implied selection for edited cells, with co-occurring neutral edits.

In sum, these data represent a comprehensive profile of base editing outcomes at endogenous DNA loci, and demonstrates the predictability and precision with which functional variants can be installed. We note that the specificity of editing is retained under strong positive selection pressure, which may be an advantage of transient expression of base editors from a doxycycline-inducible system.

### Classified *JAK1* missense mutations alter sensitivity to autologous anti-tumour T cells in primary human tumour organoids

Insensitivity to IFN*γ* in cancer cell lines is associated with inactivating mutations in the IFN*γ* pathway^3, 6^. To understand the broader functional implications of base editing variants, we mined an extensive collection of cancer cell models (*n =* 1,357) with associated exome sequencing data^33^ for pre-existing alterations at discovered *JAK1* LOF and GOF variants. The AML cell line OCI-M1 harboured the *JAK1* Val658Phe GOF mutation, and 10/1,357 cell lines had homozygous inactivating frameshift or nonsense *JAK1* mutations.

HT55 (CRC) and K2 (melanoma) cell lines harboured homozygous Glu1051Gln and Ala760Val putative *JAK1* LOF missense mutations, respectively (Supplementary Fig. 7a). As predicted, HT55 and K2 failed to respond to IFN*γ* compared to *JAK1* WT cancer cell lines, as measured by failure to induce MHC-I and PD-L1 expression (Fig. 6a). The endogenous C->T mutation in K2 cells was amenable to correction by adenine base editing. ABE8e-NGN-mediated reversion of this *JAK1* mutation led to restoration of response to IFN*γ* (Supplementary Fig. 7b), verifying that this variant is responsible for resistance to IFN*γ*. These data indicate that our base editing variant map is of broad utility, and not private to a particular cell model or tissue type. Interestingly, most of these cancer cell lines were derived before ICB was widely available, which suggests these variants arose from *in vivo* immunoediting^3, 9^ rather than acquired therapy resistance.

**Figure 6.**
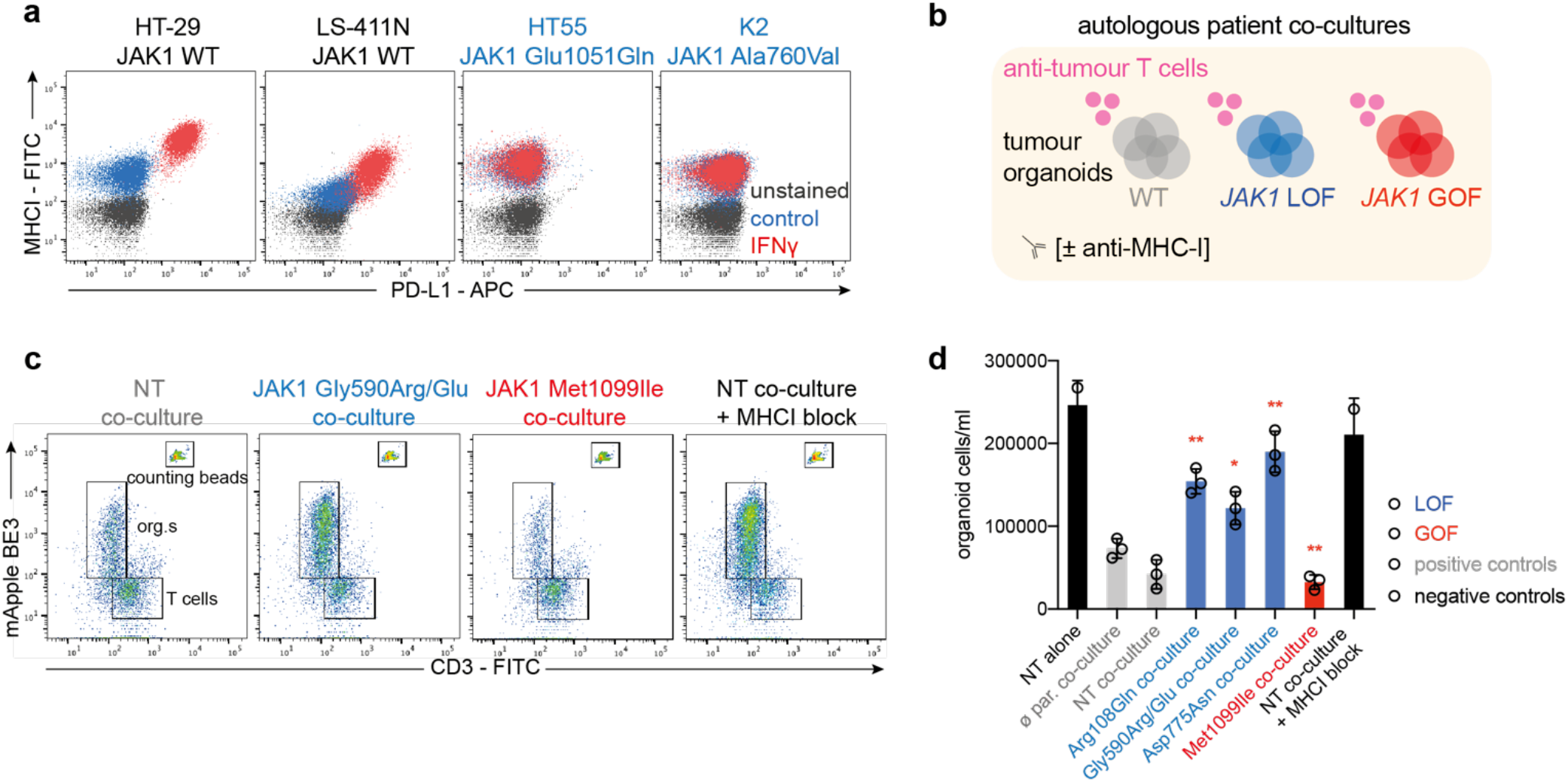
Classified *JAK1* missense mutations alter tumour organoid sensitivity to autologous anti-tumour T cells. **a)** Flow cytometry analysis of PD-L1 and MHC-I expression, showing cancer cell lines with endogenous LOF mutations in *JAK1* failing to respond to IFN*γ*. Data are representative of two independent experiments performed on separate days. **b)** Schematic of co-culture experiments to assess T cell-mediated killing of patient-derived, autologous tumour organoids, CRC-9. **c)** T cell-mediated killing of autologous human tumour organoids. Flow cytometry analysis of T cells and organoids (expressing iBE3-mApple) after 72 h of co-culture. *JAK1* base edits are indicated. Counting beads were used to quantify absolute cell counts. Data are representative of three biological replicates. **d)** Quantification of T cell-mediated killing of autologous tumour organoids from flow cytometry analysis. Data represent the average ± SD of three biological replicates, and were compared against parental co-culture controls using an unpaired, two-tailed Student’s t-test (*P* ** <0.01, *<0.05). NT, non-targeting gRNA; ø par., parental tumour organoid.

To assess the relevance of our findings in a more translational setting, we applied base editing to a primary tumour organoid (CRC-9, harbouring *FBXW7* and *TP53* driver mutations) derived from an MSI colorectal cancer patient where autologous, tumour-reactive T cells have been derived from the patient’s PBMCs^59, 60^ (Fig. 6b). Following enrichment for tumour reactive populations and expansion, co-cultured PBMCs were exclusively CD3^+^, implying a high proportion of T cells^59, 60^ (Supplementary Fig. 7c). Firstly, we confirmed that base editing of *JAK1* to install clinically observed missense variants in CRC-9 tumour organoids altered sensitivity to IFN*γ*, as measured by cell proliferation in 3D, with LOF mutations conferring resistance and the GOF mutation *JAK1* Met1099Ile increasing sensitivity (Supplementary Fig. 7c). Next, we used a co-culture of matched tumour-reactive T cells with genetically engineered tumour organoids to assess T cell mediated killing by flow cytometry (Fig. 6c). In this setting, T cell mediated killing of organoids was dependent on MHC-I, pre-exposure of organoids to IFN*γ* to increase MHC-I expression and antigen presentation, but not PD-1 inhibition with nivolumab, or CD28 co-stimulation (Supplementary Fig. 7d). Strikingly, all *JAK1* LOF mutant tumour organoids had significant resistance to anti-tumour T cell mediated killing relative to WT controls, with some mutants achieving survival comparable to antibody blockade of MHC-I, or growing tumour organoids in the absence of T cells (Fig. 6d). Conversely, the GOF mutant Met1099Ile had increased sensitivity to T cell mediated attack.

Taken together, these data illustrate that IFN*γ*-pathway variant maps from base editing screens may be prognostic of anti-tumour immunity. Our data also highlights that *JAK1* GOF can sensitise immuno-resistant *FBXW7*-mutant cancers^32^ to T cells.

## Discussion

In this report, we perform a total of 18 screens with CRISPR-Cas9 and base editors to systematically catalogue the genetic dependencies of IFN*γ* response in CRC cells, and map > 300 missense mutations affecting IFN*γ* pathway activity (see also Supplementary Discussion). Through the use of multiple cytidine and adenine base editors, to the best of our knowledge, this study represents one of the most saturating base editing mutagenesis screens performed to date^19, 20, 61^. Furthermore, we deploy base editors to systematically study protein structure and function throughout a signalling pathway. We provide BE-view as an online resource to facilitate exploration of these data: www.sanger.ac.uk/tool/be-view.

Tumour cell sensitivity to IFN*γ* is an important determinant of ICB response in multiple tumour types^5–8^. *JAK1* is mutated in approximately 10 % of CRC and 6 % of skin cutaneous melanoma, with a significant decrease in survival for melanoma patients with deleterious *JAK1* alterations^6^. We detected known LOF variants (JAK1 Asp775Asn, Trp690*)^6^ and assigned LOF to VUS in *JAK1* that may have contributed to primary or acquired resistance to ICB resistance in the clinic (e.g. *JAK1* Gly590Arg, Gly182Glu, Gly655Asp, Pro674Ser)^54, 56^. We also discovered a splice mutation in *JAK1* as a high-confidence LOF variant (Arg110 splice variant), however this LOF mutation was recorded in a patient’s tumour with a partial response to anti-CTLA-4^54^. This highlights that the presence or absence of LOF variants in the IFN*γ* pathway in a tumour biopsy is not an absolute determinant of ICB response; rather, outcome is dependent on multiple factors including the penetrance of the mutation itself (i.e. zygosity), tumour clonal architecture, co-occurring mutations, tumour mutational burden, oncogenic signalling, tumour microenvironment, antigen presentation and immune checkpoint engagement^4, 11^. Further work is required to establish the relative importance of each of these determinants, which will be increasingly feasible as the number of tumour sequencing studies increases, and as more datasets become available from matched tumour samples before and after ICB therapy. The variant database provided here will improve the interpretation of such data by enabling functional annotation of clinical variants.

IFN*γ* signalling through the JAK-STAT pathway is not only relevant for cancer immunotherapy, but also underpins pathology in myeloproliferative neoplasms, chronic mucocutaneous candidiasis, primary immunodeficiency and several inflammatory diseases^1, 2^. The molecular understanding of JAK-STAT signalling to date has been hindered by the lack of a full-length crystal structure of JAK1, and the complex intra-molecular regulation by the *JAK1* pseudokinase domain^1^. We report base editing screens mapping LOF and GOF variants in key regulatory regions of the *JAK1* pseudokinase-kinase domain interface, and conformational inter-molecular protein-protein interactions with SOCS1, demonstrating that base editing may be harnessed to understand complex protein biology, and potentially direct drug discovery efforts without prior detailed structural information.

Most of the functional variants discovered through base editing had clinical precedence (Supplementary Table 5), implying that immunoediting in cancer may be more prevalent than previously thought^3^. It is evident from this study and SGE experiments that mutation of key residues to any alternative residue can be deleterious^15^ or confer drug resistance^12^ in some contexts. Coordinated, international efforts to catalogue variant effects to understand gene function and disease have been initiated (DOI: 10.5281/zenodo.4989960). Our data highlight the exciting potential of semi-saturating base editing mutagenesis, which we envisage will complement SGE^15^, *in silico*^62^, and prime editing^63^ technologies in establishing the functional consequence of genetic variation.

## Methods

### Cell lines and culture

All cell lines were mycoplasma tested and verified by STR profiling. Cells were maintained in a 5 % CO_2_, 95 % air, humidified incubator at 37 °C, in RPMI supplemented with 1X GlutaMAX, 1X penicillin-streptomycin and 10 % FCS (Thermo Fisher Scientific). Where indicated, CellTiter-Glo proliferation assays (Promega) were performed to assess drug response following manufacturer’s instructions.

### Molecular biology and cloning

BE-FLARE reporter was synthesised as a gblock (IDT), essentially as described^39^ except where His66 codon was changed from CAC to CAT such that a single base edit can convert BFP to GFP. The gblock was integrated into a Kpn I-Eco RI digested pKLV2-gRNA expression lentiviral plasmid by Gibson assembly (NEB), expressing a BE-FLARE gRNA (5’-GCTCATGGGGTGCAGTGCTT-3’).

For generation of doxycycline-inducible base editing plasmids, we digested CLYBL-hNGN2-BSD-mApple^64^ with BamHI and PmeI (thus removing hNGN2; Addgene plasmid #124229) as a backbone and used Gibson assembly to insert PCR derived fragments containing BE3^39^, YE1-BE4max-Cas9NGN (Addgene plasmid #138159), BE3.9max-Cas9NGN^20, 44^ and ABE8e-Cas9NGN^48^.

To generate N-terminally-tagged, human, HA-SOCS1 and FLAG-JAK1 or FLAG-JAK1Gly590Arg mutant constructs, we used Addgene Plasmid #48140 as a transient expression vector backbone by removing Cas9 and GFP with EcoRI-AgeI digestion (NEB), and inserting three overlapping gBlock dsDNA fragments for JAK1, or one gBlock for SOCS1 (IDT) by Gibson assembly (NEB).

All plasmid inserts were fully sequence verified by Sanger sequencing (Eurofins). Plasmids from this article will be available from Addgene following publication (Supplementary Table 6).

### Base editor cell line generation

We knocked in base editing machinery by co-transfecting (FuGENE HD; Promega) with a plasmid encoding Cas9 and a gRNA targeting the human CLYBL locus (5’-ATGTTGGAAGGATGAGGAAA-3’), and a plasmid encoding the tet-ON base editor, blasticidin resistance and mApple expression cassettes within CLYBL homology arms. HR rates were increased by overnight pre-incubation of the cells with DNA-PK inhibitor (1 μM AZD7648). We selected transfected cells in blasticidin (10 μg/ml; Thermo Fisher Scientific) for four days and then maintained cells in 5 μg/ml thereafter. Pools were further selected by FACS for mApple expression (all positive cells). Base editing efficiency was tested using BE-FLARE^39^.

For BE3.9max NGN and ABE8e NGN, clonal lines were used for screening, which were assessed for editing activity using BE-FLARE^39^ (CBE) or a stop codon GFP reporter^65^ (ABE).

*MLH1* KO cell line clones were generated by transient transfection of a Cas9-T2A-EGFP expression plasmid (Addgene Plasmid #48140), with co-expression of an *MLH1*-targeting gRNA (5’-GCACATCGAGAGCAAGCTCC-3’), which was introduced by Golden Gate into the Bbs I site of the same plasmid. Single transfected cells were selected by EGFP expression by FACS into 96 well plates for screening by PCR and Western blotting.

### Library production

gRNAs were designed using the Wellcome Sanger Institute Genome Editing (WGE) tool^40^ https://wge.stemcell.sanger.ac.uk. Stop-essential base editing gRNA controls were selected from the iSTOP database^66^. ssDNA oligonucleotide libraries (Twist Biosciences) were resuspended and PCR amplified (KAPA HiFi HotStart ReadyMix; Roche) for 10 cycles with the addition of Gibson homology arms in the primer sequences. After PCR purification (AMPure XP SPRI beads; Beckman Coulter), we performed Gibson assembly (NEB) reactions at a 5:1 insert to vector ratio, with a BbsI-digested pKLV2-BFP-Puro lentiviral hU6 gRNA expression vector as the recipient vector^23, 67^ (Addgene Plasmid #67974). After ethanol precipitation, we performed multiple electroporations (ElectroMAX Stbl4 cells; Thermo Fisher Scientific) to maintain library complexity. Transformation efficiency was verified by serial dilution of the liquid culture onto LB+Amp agar plates. Library plasmid pools were propagated in liquid culture in LB with ampicillin (100 μg/ml) at 30 °C overnight and extracted (Qiagen).

HEK293T cells were co-transfected with psPAX2, pMD2.G and the lentiviral gRNA plasmid at a 3:1:5 mass ratio using FuGENE HD (Promega) in Opti-MEM (Thermo Fisher Scientific). Media was refreshed the next day and viral supernatant was harvested 72 h post-transfection, filtered and frozen. Thawed viral supernatant titre was assessed by infection of HT-29 cells, always in the presence of 8 μg/ml polybrene (Sigma-Aldrich), and 48 h later, measuring BFP expression by flow cytometry.

### CRISPR-Cas9 KO screens

A custom gRNA library was manually designed from an extensive literature search, generated (Oxford Genetics), titrated using an mCherry fluorophore, and used at a viral titre that achieved 30-50 % infection in HT-29 and LS-411N cells stably expressing Cas9^23^. Cells were selected with puromycin for 4 d (2 μg/ml and 1 μg/ml, respectively), maintaining 300 X coverage, with a time 0 (T0) control sample taken 7 d after infection. 10 d after infection, cells were selected with IFN*γ* (2000 U/ml; Thermo Fisher Scientific) for a total of 7 d with the IFN*γ* arm having IFN*γ* media refreshed after 4 d and the control arm being passaged after 4 d. Each screen was performed independently twice on separate days.

### Base editing screens

Base editing screens were performed with a gRNA coverage of 400-1000-fold. We adopted viral doses achieving 30-50 % infected cells. For proliferation screens, as with the CRISPR-Cas9 KO screens described above, we selected cells for 4 d with puromycin, a T0 sample was taken at 6 d post-infection, then doxycycline (1 μg/ml) was added for 3 d to induce base editing, followed by selection with IFN*γ* (2000 U/ml; Thermo Fisher Scientific) for 7 d. For the FACS screens, the library-transduced and puromycin-selected cell population was base edited by the addition of doxycycline 10 d after infection for 3 d, and 14 d after infection, IFN*γ* (400 U/ml) was added to induce PD-L1 and MHC-I expression for 48 h before FACS. Due to lower overall editing efficiencies for BE4max-YE1 compared to BE3, we extended the selection with IFN*γ* from 7 days to 14 days and did not perform a FACS selection assay to maintain good library representation. All screens were performed independently twice on separate days.

### FACS and flow cytometry analysis

Cells were washed were harvested, washed once in FACS buffer (0.5 % FCS, 2 mM EDTA in PBS) before staining on ice for 30 min in the dark with anti-PD-L1 (MIH1; APC) and anti-MHC-I (W6/32; FITC; both 1:100 dilution; Thermo Fisher Scientific), and washed twice in FACS buffer and adding DAPI (1 μg/ml; Thermo Fisher Scientific) before analysis (LSRFortessa; BD Biosciences). For base editing screens, FACS was used to sort approximately 250,000 LOF cells (BD Influx cell sorter; BD Biosciences), which were expanded for seven days in the absence of IFN*γ* before DNA extraction. For experiments with HT55 and K2, cells were treated with IFN*γ* (400 U/ml; Thermo Fisher Scientific) for 48 h before analysis. FACS data were analysed with FlowJo software. For *JAK1* variant SNP correction in K2 cells, we generated ABE83-NGN doxycycline-inducible derivative and introduced the lentiviral gRNAs 5’-GAGGAACAATCCATGGGATT-3’ (JAK1) or 5’-GCTGATATATACGACAAGCC-3’ (NT control), as described above. Three days after addition of doxycycline (1 μg/ml), we stimulated the cells with IFN*γ* (400 U/ml; Thermo Fisher Scientific) for 48 h before flow cytometry analysis.

### Next generation sequencing

Amplicon sequencing was performed as described^68^ with primers provided in Supplementary Table 7. Amplicons for *JAK1* 5’UTR and 724 positions failed quality control. For gRNA sequencing, genomic DNA was extracted from cell pellets from CRISPR or base editing screens (DNeasy Blood & Tissue; Qiagen), gRNA DNA sequences were PCR amplified (empirically determined number of cycles; KAPA HiFi HotStart ReadyMix; Roche), SPRI purified (AMPure XP SPRI beads; Beckman Coulter) and quantified (Qubit; Thermo Fisher Scientific). In addition, plasmid DNA from the original library always served as a control in screening experiments. PCR products were then indexed with a second round of PCR (8-10 cycles) with unique identifier sequences and Illumina adapters, SPRI-purified, quantified (Bioanalyzer; Agilent), pooled in an equimolar ratio, quantified by qPCR and sequenced on a HiSeq2500 (Illumina) with a custom sequencing primer (5’-TCTTCCGATCTCTTGTGGAAAGGACGAAACACCG-3’) for 19 bp single-end reads of the gRNA on Rapid Run Mode.

### Validation experiments

Individual gRNAs were cloned in an arrayed format using a Golden Gate-based approach. We designed primers encoding a gRNA with BbsI overhangs and an additional G for hU6 RNApolIII transcription (Forward: 5’-CACC**G**NNNNNNNNNNNNNNNNNNN-3’ and Reverse: 5’-AAACNNNNNNNNNNNNNNNNNN**C**-3’), annealed by boiling and slowly cooling to room temperature, and then ligated duplexes with a BbsI entry vector, BbsI-HF (NEB), T4 DNA ligase and buffer (NEB), 1X BSA (NEB) for 30X cutting (37 °C) and ligating (16 °C) cycles, before heat-shock transformation of DH5-*α E. coli* (NEB).

### Western blotting

Cells were lysed with 4X sample loading buffer (8% SDS, 20 % *β*-mercaptoethanol, 40 % glycerol, 0.01 % bromophenol blue, 0.2 M Tris-HCL pH 6.8) supplemented with benzonase (Sigma) to digest genomic DNA. Samples were boiled for 5 min at 95 °C before SDS PAGE (Thermo Fisher Scientific). PVDF membranes were probed with the following primary antibodies: STAT1 (#9172S), *JAK1* (#50996), p-STAT1 (#9167), *β*-tubulin (#2146) (Cell Signaling Technology). Secondary antibodies were conjugated to horseradish peroxidase. For validation experiments, LOF mutant *JAK1* edited cells were pre-selected with IFN*γ* for 5 days prior to re-stimulation to enrich for edited cells. These experiments were performed without pre-selection with similar results but smaller differences due to the presence of unedited cells. For stimulation of JAK-STAT signalling, cells were treated with IFN*γ* (400 U/ml; Thermo Fisher Scientific) for 1 h.

### Immunoprecipitation

HEK293T cells were transfected with FLAG-JAK1 or FLAG-JAK1Gly590Arg and HA-SOCS1 (FuGENE HD, Promega). 72 h later, cells were stimulated with IFN*γ* (400 U/ml; Thermo Fisher Scientific), or RPMI complete medium as a control, for 1 h before lysis with lysis buffer (20 mM Tris-HCl pH 7.4, 137.5 mM NaCl, 10 % glycerol, 1 % Triton X-100) supplemented with benzonase and protease-phosphatase inhibitor cocktail (Sigma-Aldrich). 25 ul of protein G Dynabeads (Thermo Fisher Scientific) were conjugated to 1 μg of anti-FLAG antibody (M2; Sigma-Aldrich) for each immunoprecipitation, which was carried out overnight at 4 °C with inversion. The following day, beads were washed with wash buffer (lysis buffer with 0.1 % Triton X-100), before elution with 4X sample loading buffer and SDS-PAGE. We used beads alone (without anti-FLAG antibody) as a control for binding specificity.

### Data analysis

To call SNPs from amplicon sequencing, we used CaVEMan^69^ and BCFtools^70^. Variant allele frequency (VAF) was calculated using vafCorrect^71^, and variants with <1 % VAF were filtered out. For COSMIC analysis, mutations and frequencies were downloaded from https://cancer.sanger.ac.uk/cosmic in January 2021. For visualisation of crystal structures, we used PyMOL (version 2.4.1), for graphs we used GraphPad Prism (version 8) or R ggplot2 (3.3.0). For CRISPR-Cas9 and base editing screens, we filtered out any gRNAs with 0 read counts in the control samples. Log2 fold-changes (L2FC) were calculated from normalised read counts (normalised reads per million = gRNA reads/total reads for the sample x 1,000,000 + 1 pseudocount). For CRISPR-Cas9 screens, MAGeCK analysis was performed using default parameters, except that normalization is set to ‘none’, as the input corrected counts had already been normalised. A false discovery rate cut-off of 5% (FDR ≤ 0.05) was applied to identify the significant genes. For base editing screens, we implemented DrugZ to calculate a gene level z-score for each fold change using an empirical Bayes estimate of the standard deviation. We calculated z-scores using normalisation by L2FC from nonessential/intergenic/non-targeting control gRNAs. Analyses with L2FC and z-scores gave similar results. For base editing screens, we considered the base edits from each gRNA as single mutations or the mutation of all cytosines or adenines in the base editing window and used VEP^72^ to assign amino acid changes. For BE3 NGG we assumed a lenient window of 4-9 and for BE4max-YE1 NGN we used a window of 5-7, where 20-23 is the PAM. We focussed our analysis on VEP output of MAINE selected canonical protein coding transcripts. For annotation of edit consequence, we consolidated multiple predicted consequences by giving priority to the most deleterious as follows: stop gain > start loss > splice variant > missense > UTR > synonymous variant. For base editing screens, we filtered out samples with < 100 gRNA read counts for any sample in either replicate, and one gRNA that was over-represented (> 50,000 reads) in the library. Data wrangling for graphs was performed with R and can be found here: https://github.com/MatthewACoelho/Base_Editing_Screens.

### Data availability

All sequencing data have been released to the European Nucleotide Archive (ENA) for public access (Supplementary Table 8). Read counts for CRISPR and base editing screens are available as Supplementary Tables.

### qPCR

72 h after base editing (induced by the addition of doxycycline), RNA was extracted and genomic DNA was removed (RNeasy columns and DNase I; Qiagen), followed by cDNA synthesis with SuperScript IV and random hexamers, and analysis using SYBR Green reagents on the Step One Plus (Thermo Fisher Scientific), with the following primers: Human *JAK1* 5’-GAGACAGGTCTCCCACAAACAC-3’, 5’-GTGGTAAGGACATCGCTTTTCCG-3’, Human *GAPDH* 5’-GTCTCCTCTGACTTCAACAGCG-3’, 5’-ACCACCCTGTTGCTGTAGCCAA-3’.

### Giemsa staining

After six days of selection with IFN*γ* (1500 U/ml; Thermo Fisher Scientific), cells were washed with PBS, fixed with 4 % PFA for 20 min and then stained with Giemsa working solution (1X in water; Sigma-Aldrich) for 2 h at room temperature with gentle rocking. Wells were rinsed with deionised water three times and then allowed to dry before images were taken by scanning.

### Co-cultures with autologous T cells

Derivation of tumour organoids, enrichment of tumour reactive T cell populations from patient PBMCs and co-culture killing assays were performed as described^59^. Briefly, CRC-9 cells were pre-stimulated with IFN*γ* (Thermo Fisher Scientific, 400 U/ml) overnight to increase MHC-I expression, then seeded in suspension in non-tissue culture treated 96 well plates at a 3:1 E:T ratio for 72 h, with or without anti-CD-28 coating, nivolumab (20 ug/ml; Selleckchem), and MHC-I blocking antibody (W6/32; 50 ug/ml) in RPMI supplemented with human serum and primocin (Invivogen). Cells were harvested and stained with anti-CD3 FITC antibody (UCHT1; Thermo Fisher Scientific, 1:100), washed in FACS buffer before the addition of DAPI and flow cytometry analysis. 123count eBead counting beads (Thermo Fisher Scientific) allowed for quantification of absolute cell counts based on volumetric measurements from bead counts. Growth of organoids in 3D was achieved by growth in 80 % basement membrane extract (BME; R&D Systems).

## Acknowledgements

We thank the Translational Cancer Genomics team and iFuncOnc project team, especially Gabriele Picco, for helpful discussion, and Katrina McCarten for critical reading of the manuscript. We thank Sanger DNA pipelines and Mamta Sharma for assistance. We thank Nick Boughton for assistance with BE-view website support.

## Author Contributions

MAC, AB, EV and MJG devised the study. MAC carried out the experiments and MAC, EK and SB performed data analysis. TB, SC, EG, CD, SC, JR, MT and MAC designed and produced gRNA libraries. FG, SV, and CH assisted with experiments and sequencing. SB and EV, SC, CMC and VV developed and provided essential reagents. MAC and MJG wrote and all authors reviewed the manuscript.

## Supplementary References

**Supplementary Figure 1.**
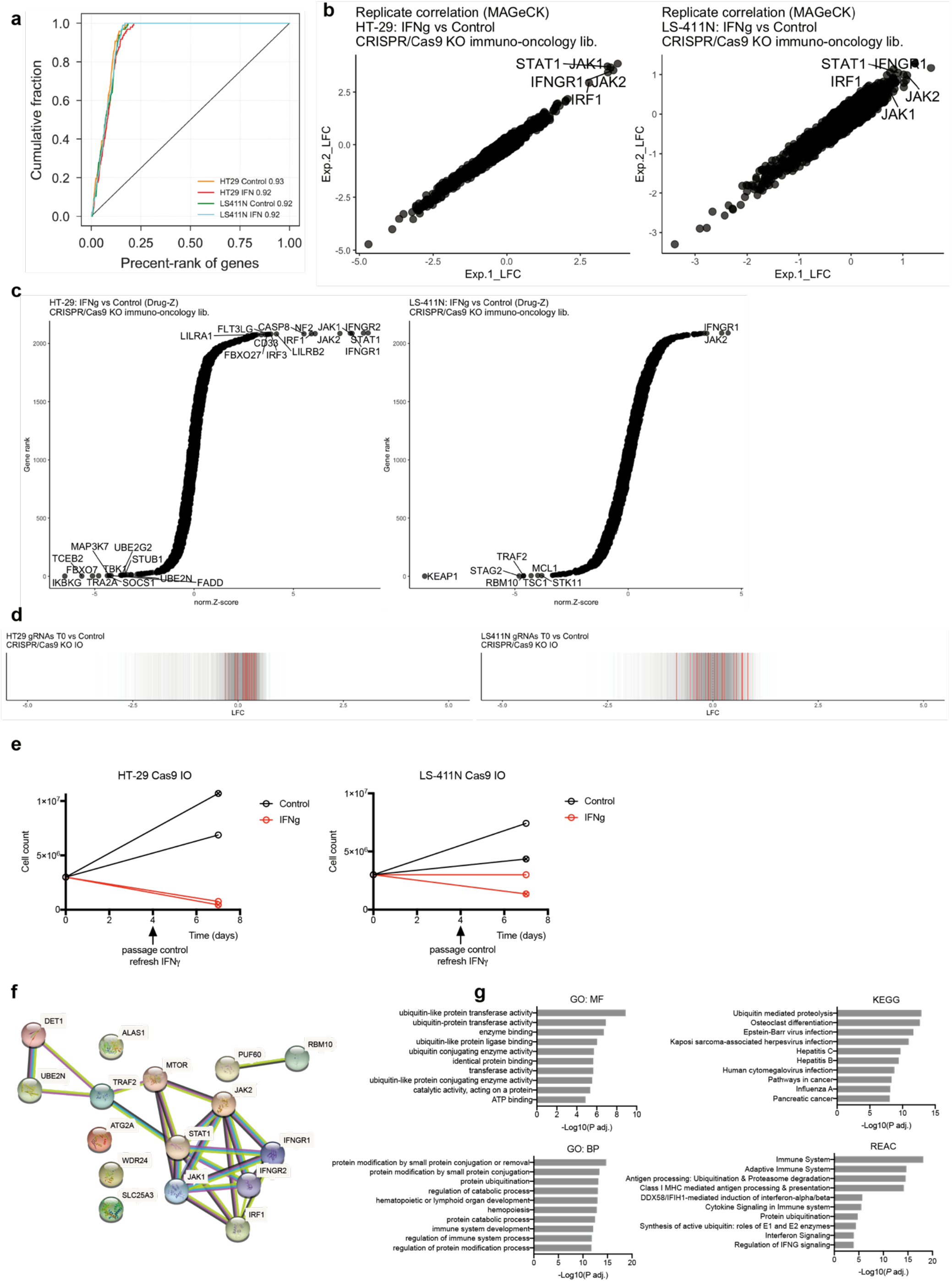
CRISPR/Cas9 screens identify mediators of IFNγ sensitivity and resistance. **a)** Precision-recall analysis of CRISPR/Cas9 screen performance in HT-29, LS-411N cells with or without IFN*γ*. Precision-recall was based on the recovery of known essential genes versus the plasmid control, and the area under the curve is given in each case. **b)** Replicate correlation from MAGeCK analysis of CRISPR/Cas9 screens (control vs IFN*γ* arms) based on gRNA log2 fold-changes. Top resistance hits are shown for each cell line. **c)** Drug-Z analysis of averaged CRISPR/Cas9 screens (control vs IFN*γ* arms) with top hits indicated for each cell line. **d)** MAGeCK analysis of CRISPR/Cas9 screens (control vs T0 arms) showing individual gRNAs targeting *JAK1*, *JAK2*, *IFNGR1*, *IFNGR2*, *STAT1*, *IRF1*, in red. **e)** Growth curves showing cell proliferation in two independent CRISPR/Cas9 immuno-oncology target screens performed in HT-29 and LS-411N CRC Cas9-expressing cell lines. Arrow indicates when the cells were passaged in the control arm, whereas at this point in the IFN*γ* arm, IFN*γ* was refreshed. **f)** STRING network analysis of protein interactions for IFN*γ*-sensitising and resistance genes common to HT-29 and LS-411N. **g)** Gene ontology analysis of shared CRISPR-Cas9 gene hits shows enrichment for genes involved in protein ubiquitination (based on molecular function “MF”, Biological Process “BP”, KEGG, and Reactome; g:profiler).

**Supplementary Figure 2.**
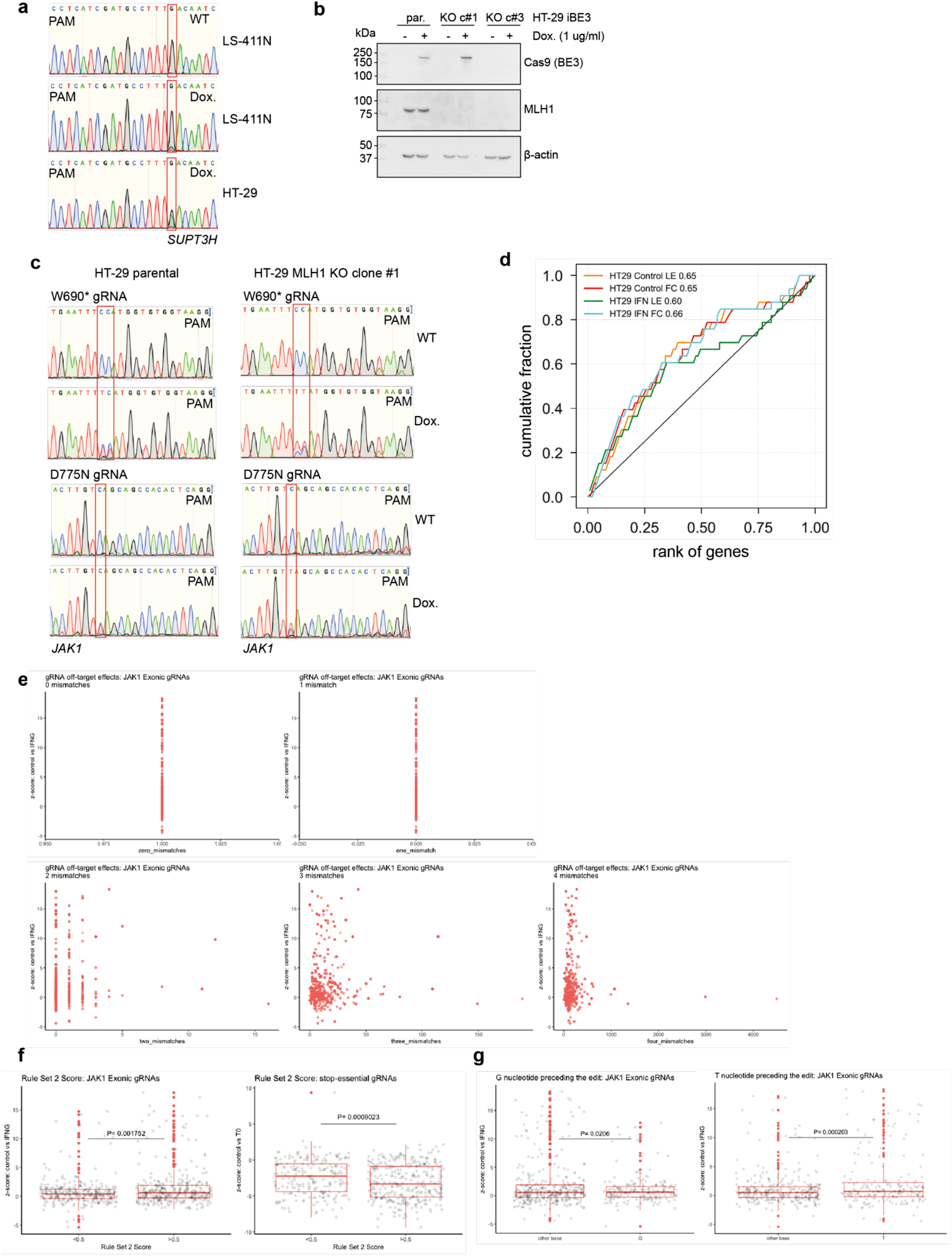
Base editing mutagenesis screening of *JAK1* variants. **a)** Sanger sequencing analysis of the *SUPT3H* locus targeted with BE3 in HT-29 and LS-411N iBE3 cells. G->A editing is observed with the addition of doxycycline for 72 h. The protospacer sequence is displayed. **b)** Western blot analysis of HT29 iBE3 *MLH1* KO single cell clone (KO c#3). KO was performed using transient expression of a CRISPR/Cas9 plasmid co-expressing a gRNA against *MLH1*. **c)** Sanger sequencing analysis of base editing of *JAK1* loci using the indicated gRNAs in HT-29 iBE3 and HT-29 iBE3 *MLH1* KO cells. Base editing was induced with doxycycline for 72 h. **d)** Precision-recall analysis of base editing screen performance in HT-29 iBE3 cells in the control or IFN*γ* arms based on the recall of known essential genes. Area under the curve is given in each case for Drug-Z analysis of average control vs time zero (T0) conditions from two independent replicate screens. (FACS screen, fc; Proliferation screen, Le). **e)** Off-target analysis of *JAK1* base editing library. Plotted are the proliferation screen z-scores (control vs IFN*γ* arms) against the number of off-target genomic positions (with 0 = on-target, 1, 2, 3 and four mismatches) for each gRNA targeting *JAK1* exonic regions. **f)** gRNAs targeting *JAK1* exons or generating stop codons in essential genes were assigned a Rule Set 2 Score and grouped into <0.5 or >0.5. Proliferation screen z-scores were compared between groups using an unpaired, two-tailed Student’s t-test. **g)** gRNAs targeting *JAK1* exons were grouped by the predicted edited cytosine’s direct genomic context; preceded by a G or preceded by a T. Proliferation screen z-scores were compared between groups using an unpaired, two-tailed Student’s t-test.

**Supplementary Figure 3.**
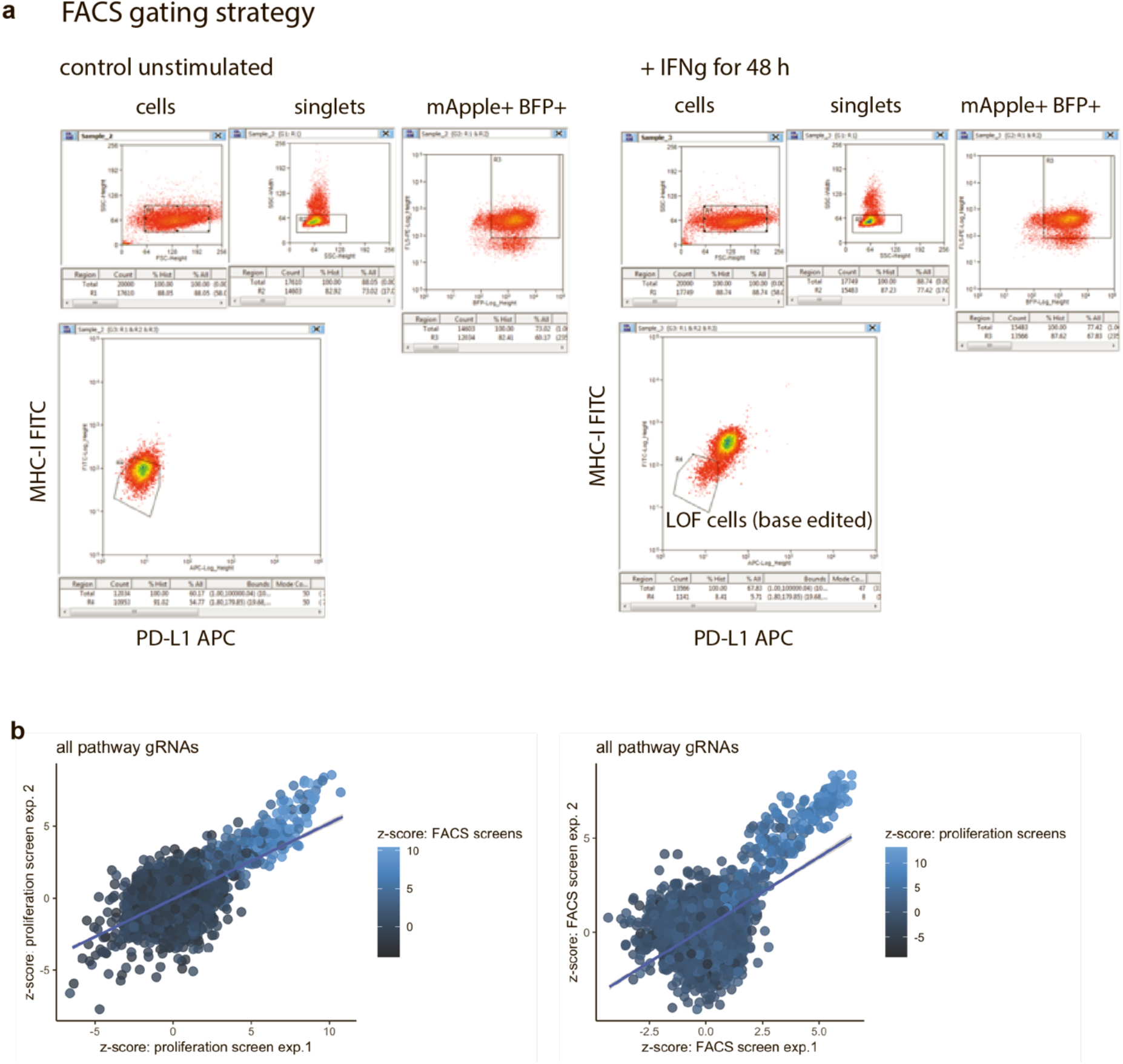
Base editing mutagenesis of the IFNγ pathway. **a)** FACS gating strategy for cells with LOF in the IFN*γ* pathway. HT-29 iBE3 cells were stimulated with IFN*γ* (400 U/ml) for 48 h before FACS. Single cells expressing base editor (mApple) and gRNA (BFP) were gated and the cells unable to induce PD-L1 and MHC-I were gated based on a unstimulated control population. Data are representative of two independent experiments performed on separate days. **b)** Replicate correlation for base editor screening of the IFN*γ* pathway. Correlation between z-scores for independent base editor screening replicate experiments performed on separate days, and independent screening assays (FACS and proliferation).

**Supplementary Figure 4.**
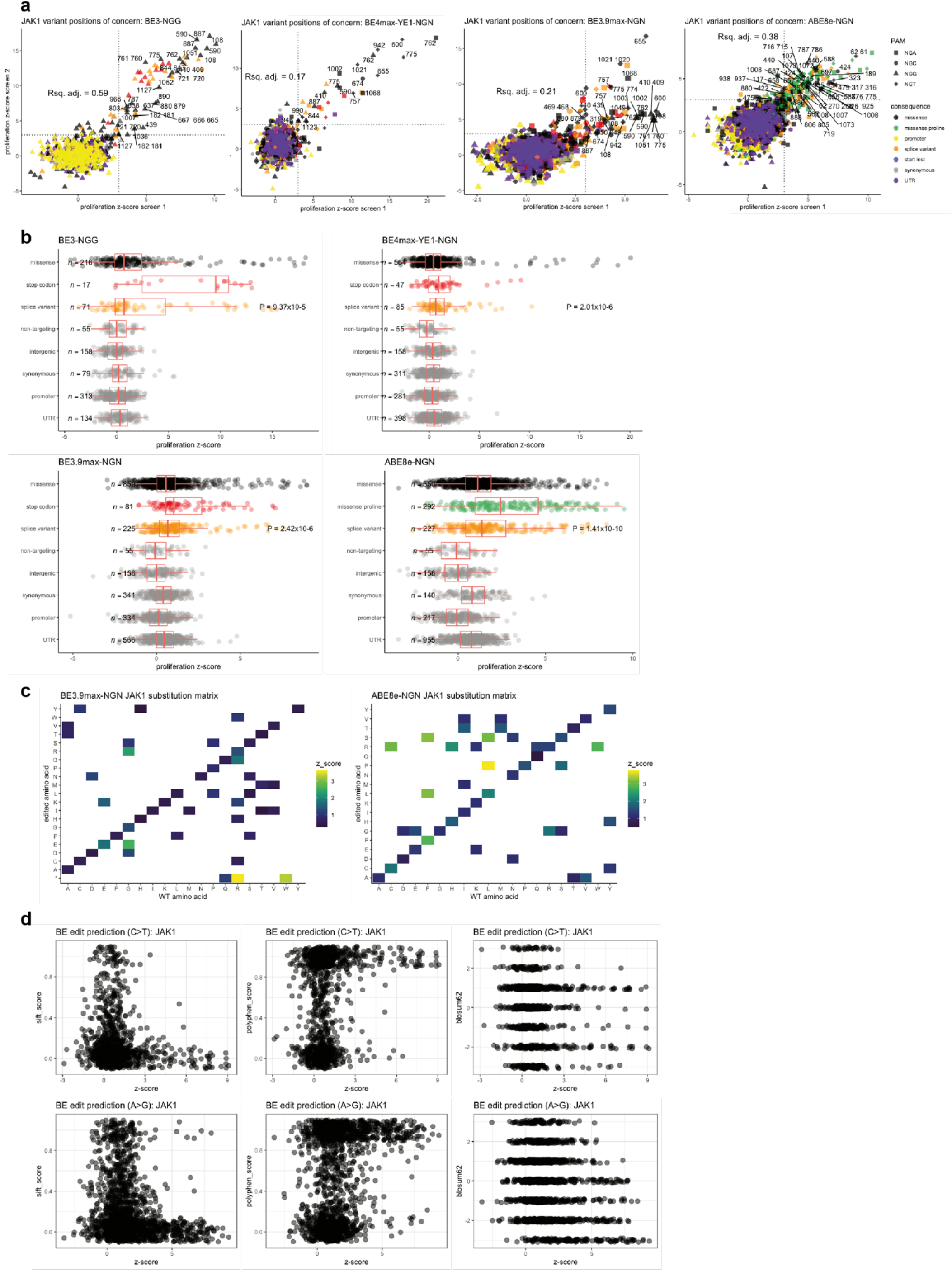
Base editing reveals *JAK1* LOF and GOF variants with clinical precedence. **a)** Replicate correlation of base editing screens using different base editor architectures and deaminases. Dot plots of gRNAs targeting *JAK1* are coloured by predicted consequence. Shape indicates PAM usage of the gRNA and adjusted R^2^ values are indicated. z-scores (control vs IFN*γ*-arms; proliferation screens) are from two independent screens performed on separate days. **b)** Boxplot of proliferation screen z-scores for gRNAs by predicted consequence. Z-scores for predicted splice variant and non-targeting gRNAs (control vs IFN*γ*-arms) were compared using an unpaired, two-tailed Student’s t-test. Shown is the median, box limits are upper and lower quartiles, whiskers are 1.5× interquartile range, and points are outliers. **c)** Heatmap amino acid substitution matrix, showing aggregated predicted codon changes for each gRNA targeting *JAK1* and gRNA z-scores from control vs IFN*γ*-arms for BE3.9max-NGN and ABE8e-NGN proliferation screens. **d)** Comparison of bioinformatic prediction of variant effect with experimental data from base editing screens (z-scores from control vs IFN*γ*-arms; proliferation screens). SIFT (0 is deleterious, 1 is tolerated), PolyPhen (0 is benign, 1 is damaging) and BLOSUM62 (positive is conserved, negative is not conserved).

**Supplementary Figure 5.**
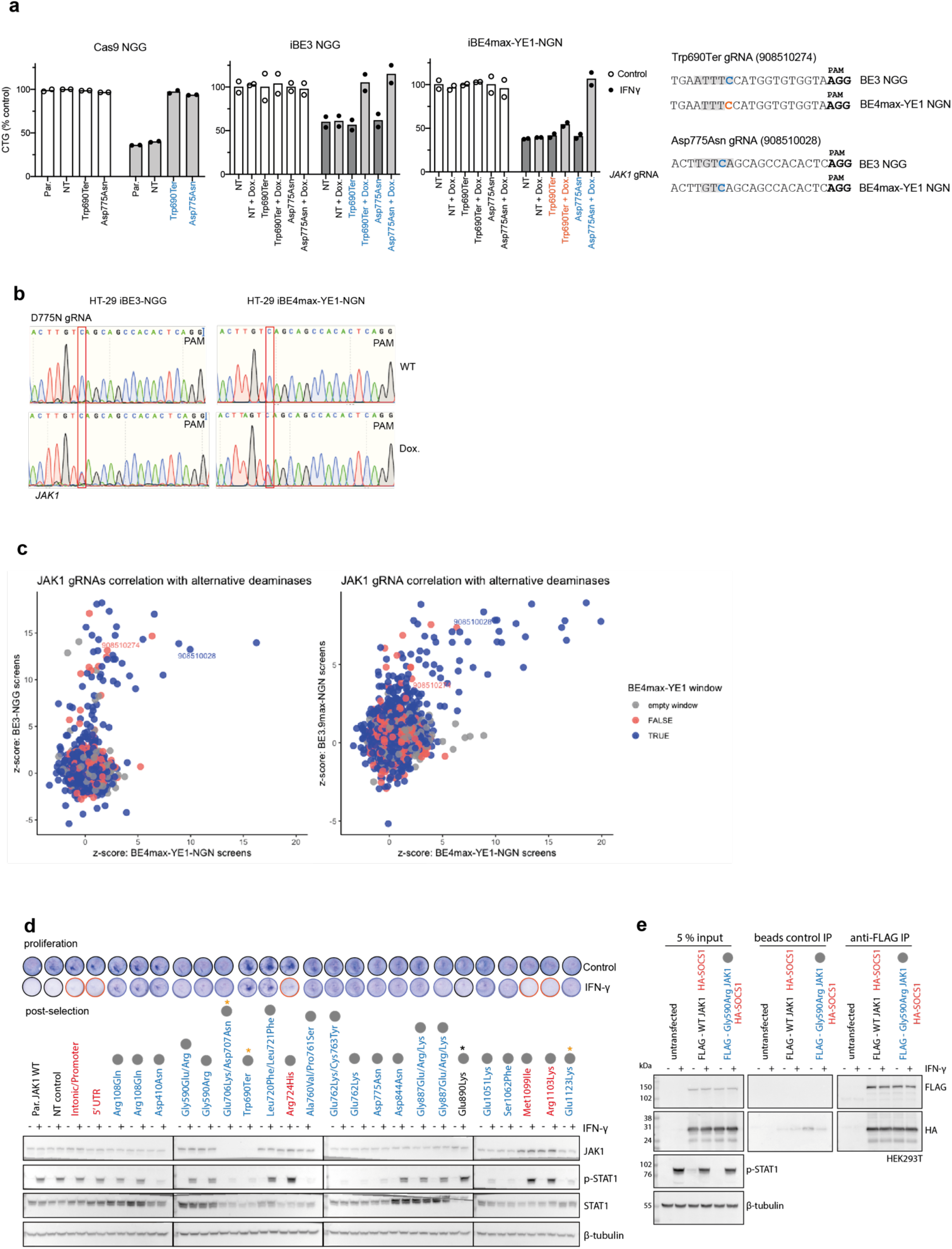
Functional validation of base editing variants conferring altered sensitivity to IFNγ. **a)** Comparison of gene editing technologies. Cas9-NGG or doxycycline-inducible BE3-NGG or BE4max-YE1-NGN were compared by measuring growth of HT-29 cells expressing the indicated gRNAs treated with IFN*γ* for 6 d. Data represent the mean of two independent experiments performed on separate days, with each experiment performed in technical triplicate. Two *JAK1* LOF gRNAs with targeted cytosines inside or outside of the predicted deaminase activity window (shaded grey). **b)** Comparison of *JAK1* base editing efficiency by BE3-NGG and BE4max-YE1-NGN. Data for HT-29 iBE3 are also shown in Supplementary Fig. 2. **c)** Correlation between gRNA performance for gRNAs in both iBE3-NGG and iBE4max-YE1-NGN, and iBE3.9-NGN and iBE4max-NGN screens. gRNAs with a target cytosine within the narrower iBE4max-YE1-NGN deaminase activity window are shown in blue. gRNA IDs relating to other Figures are shown for reference. **d)** Validation of *JAK1* variants. Independent experiments replicating phenotypes described in Fig. 4a. **e)** Immunoprecipitation analysis of HA-SOCS1 and FLAG-JAK1 or FLAG-JAK1Gly590Arg mutant from transiently transfected HEK293T cells, with and without IFN*γ* stimulation.

**Supplementary Figure 6.**
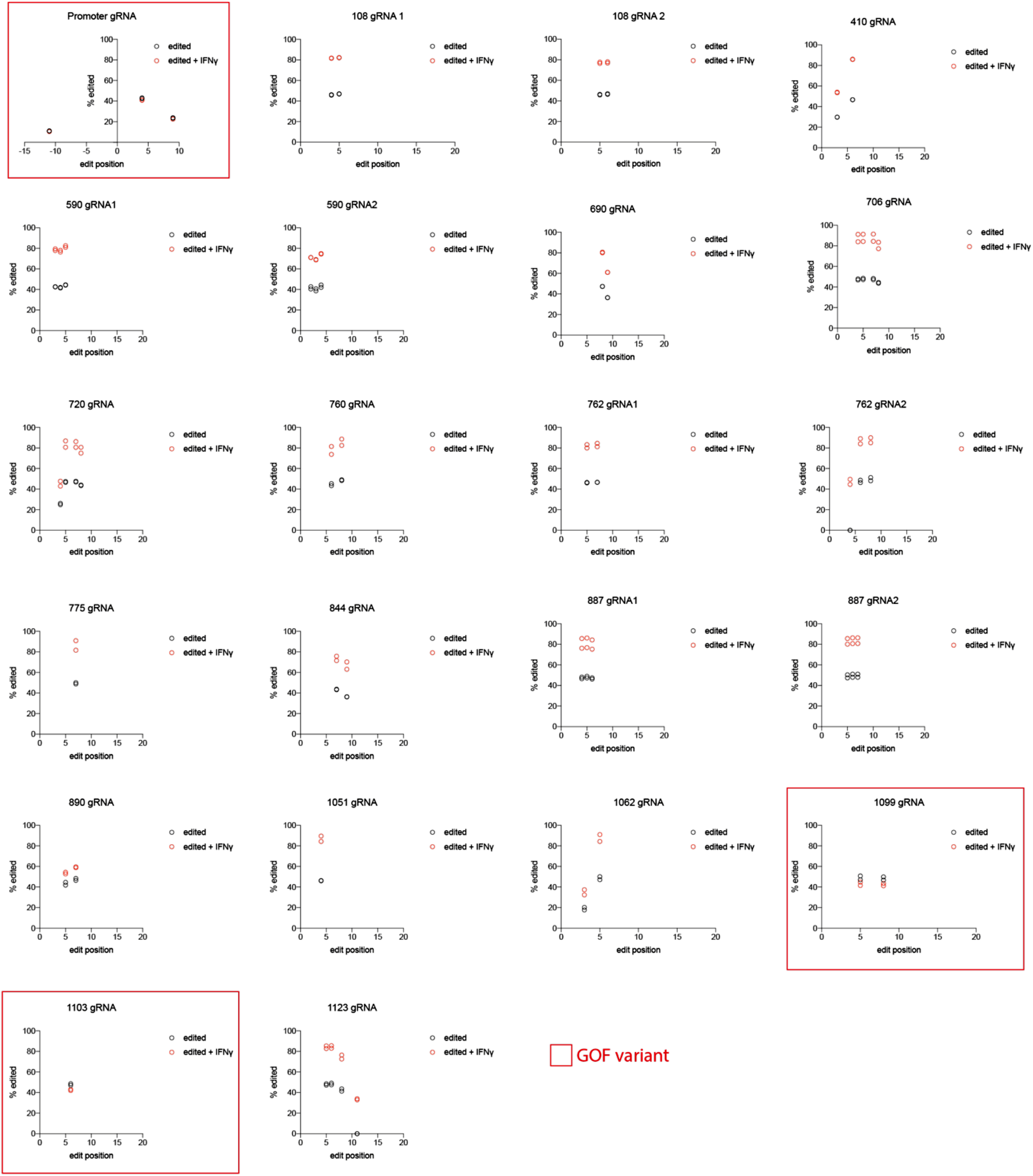
Amplicon sequencing of *JAK1* following base editing. Amplicon sequencing of endogenous *JAK1* DNA reveals the editing profile of BE3 gRNAs. Position of edits relative to the protospacer are shown for LOF and GOF gRNAs in the validation cohort. Data are generated from control cells, cells with base editing or base editing and selection with IFN*γ* for 6 d. Data represent the mean of two independent experiments performed on separate days.

**Supplementary Figure 7.**
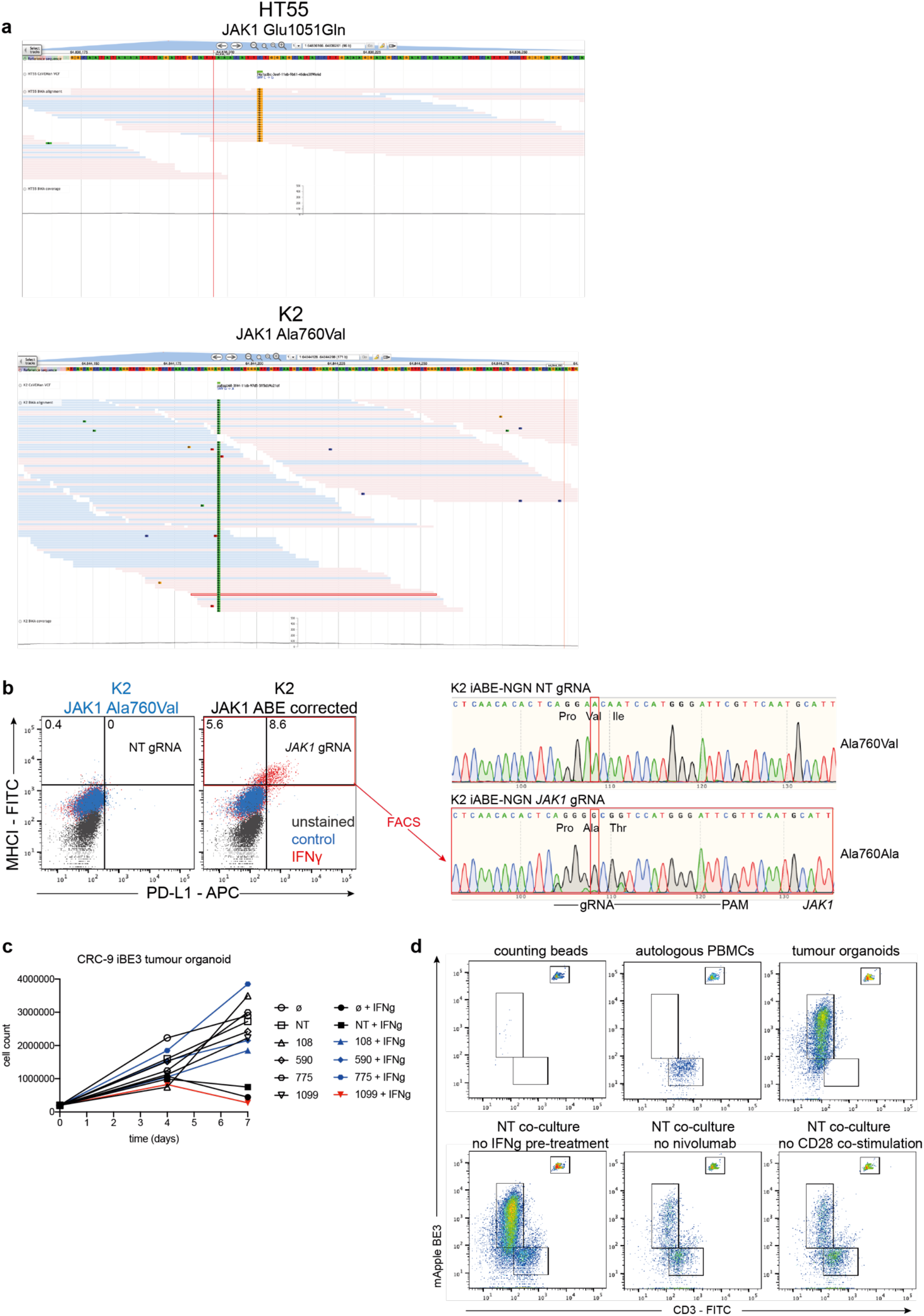
Base editing mutagenesis of the IFNγ pathway, and Classified *JAK1* missense mutations alter sensitivity to autologous anti-tumour T cells in primary human tumour organoids. **a)** Exome sequencing data from HT55 and K2 cells lines with sequencing reads showing homozygous mutations in *JAK1*. **b)** Flow cytometry analysis of PD-L1 and MHC-I expression following correction of an endogenous *JAK1* LOF mutation with ABE8e-NGN in K2 cells. Numbers represent the percentage of IFN*γ*-stimulated cells inducing expression of MHC-I and PD-L1 over baseline levels. MHC-I^+^ PD-L1^+^ cells were sorted with FACS for DNA analysis by Sanger sequencing (right panel), revealing efficient reversion to WT *JAK1* Ala760, and the bystander edit Ile759Thr. Data are representative of two independent experiments performed on separate days. NT; non-targeting control gRNA. **c)** Cell counts quantification of CRC-9 organoid growth in 3D, with (closed symbols) and without IFN*γ* (open symbols). *JAK1* LOF mutants in blue grow progressively, whereas GOF *JAK1* mutants in red, or controls in black, stop growing. Data are representative of two independent experiments performed on separate weeks. **d)** Representative flow cytometry plots and controls from the T-cell and autologous organoid co-cultures. Top panel shows counting beads, PBMCs or tumour organoids alone. Bottom panel show co-cultures after 3 d, where there is no organoids pre-treatment with IFN*γ*, no anti-PD-1 nivolumab in the co-culture, or no anti-CD28 co-stimulation. Data are representative of two-three biological replicates in each case.

## Supplementary Discussion

CRISPR-Cas9 screening identified druggable targets that sensitised tumour cells to IFN*γ* when inactivated, such as MCL1 and TBK1, highlighting potential ICB-combination therapies in CRC. In line with this, TBK1 inhibition has been reported to increase immune reactivity to tumour organoids *ex vivo*^1^. Conversely, we revealed that mTOR inactivation can facilitate tumour-intrinsic resistance to IFN*γ*, arguing against combining mTOR inhibitors with ICB in CRC^2^, although our reductionist approach does not consider the potential effects of these drugs on immune cells. Interestingly, inactivation of *KEAP1*, *FBXW7*, *NF2*, and *STK11*, modulated sensitivity to IFN*γ*, emphasising important non-cell autonomous roles for these tumour suppressor genes. KO of *NF2* resulted in increased resistance to IFN*γ*, and has also been linked to BRAF inhibitor resistance^3, 4^, consistent with an overlap between ICB resistance and MAPK inhibitor resistance pathways^5^, with possible implications for the efficacy of ICB in melanoma patients pre-treated with BRAF inhibitors.

